# Assessing the durability and efficiency of landscape-based strategies to deploy plant resistance to pathogens

**DOI:** 10.1101/260836

**Authors:** Loup Rimbaud, Julien Papaïx, Jean-François Rey, Luke G. Barrett, Peter H. Thrall

**Affiliations:** CSIRO Agriculture and Food, Canberra, ACT, Australia; BioSP, INRA, Avignon, France

**Keywords:** demo-genetic modelling, deployment strategies, durable plant resistance, pathogen evolution, quantitative resistance, rust of wheat, spatially explicit stochastic modelling, spatiotemporal simulation model

## Abstract

Genetically-controlled plant resistance can reduce the damage caused by pathogens. However, pathogens have the ability to evolve and overcome such resistance. This often occurs quickly after resistance is deployed, resulting in significant crop losses and a continuing need to develop new resistant cultivars. To tackle this issue, several strategies have been proposed to constrain the evolution of pathogen populations and thus increase genetic resistance durability. These strategies mainly rely on varying different combinations of resistance sources across time (crop rotations) and space. The spatial scale of deployment can vary from multiple resistance sources occurring in a single cultivar (pyramiding), in different cultivars within the same field (cultivar mixtures) or in different fields (mosaics). However, experimental comparison of the efficiency (i.e. ability to reduce disease impact) and durability (i.e. ability to limit pathogen evolution and delay resistance breakdown) of landscape-scale deployment strategies presents major logistical challenges.

Therefore, we developed a spatially explicit stochastic model able to assess the epidemiological and evolutionary outcomes of the four major deployment options described above, including both qualitative resistance (i.e. major genes) and quantitative resistance traits against several components of pathogen aggressiveness: infection rate, latent period duration, propagule production rate, and infectious period duration. This model, implemented in the R package *landsepi*, provides a new and useful tool to assess the performance of a wide range of deployment options, and helps investigate the effect of landscape, epidemiological and evolutionary parameters.

This article describes the model and its parameterisation for rust diseases of cereal crops, caused by fungi of the genus *Puccinia*. To illustrate the model, we use it to assess the epidemiological and evolutionary potential of the combination of a major gene and different traits of quantitative resistance. The comparison of the four major deployment strategies described above will be the objective of future studies.

**Author summary:** There are many recent examples which demonstrate the evolutionary potential of plant pathogens to overcome the resistances deployed in agricultural landscapes to protect our crops. Increasingly, it is recognised that how resistance is deployed spatially and temporally can impact on rates of pathogen evolution and resistance breakdown. Such deployment strategies are mainly based on the combination of several sources of resistance at different spatiotemporal scales. However, comparison of these strategies in a predictive sense is not an easy task, owing to the logistical difficulties associated with experiments involving the spread of a pathogen at large spatio-temporal scales. Moreover, both the durability of a strategy and the epidemiological protection it provides to crops must be assessed since these evaluation criteria are not necessarily correlated. Surprisingly, no current simulation model allows a thorough comparison of the different options. Here we describe a spatio-temporal model able to simulate a wide range of deployment strategies and resistance sources. This model, implemented in the R package *landsepi*, facilitates assessment of both epidemiological and evolutionary outcomes across simulated scenarios. In this work, the model is used to investigate the combination of different sources of resistance against fungal diseases such as rusts of cereal crops.

## Introduction

The deployment of resistant cultivars in agricultural landscapes aims to reduce the ability of plant pathogens to cause disease on crops. However, the durability of plant resistance has often been limited by evolutionary changes in pathogen populations [1]. Typically, there are two main types of resistance. Although exceptions exist, qualitative (or ‘major gene’) resistance is usually monogenic and complete, i.e. conferring full immunity [2–4]. In contrast, quantitative resistance is mostly polygenic and partial, i.e. infection is still possible but pathogen development is reduced to a greater or lesser extent. Consequently, quantitative resistance is often described as affecting one or more components of pathogen aggressiveness (defined as the quantitative ability to colonise and cause damage to the host): lower rate of infection, longer latent period, reduced propagule production, shorter infectious period or lower toxin production [5–8].

Regardless of the source of resistance, pathogens have the potential to evolve infectivity (defined as the qualitative ability to infect the host) and aggressiveness in response to the selection posed by plant resistance [4, 9]. This adaptation of pathogens at the population level is generally driven by mutations from non-adapted pathogen genotypes, changes in frequencies of pre-existing adapted genotypes, or introductions via migration from distant areas [9–11]. Emergence of novel pathotypes can result in the breakdown of qualitative resistance or the erosion of quantitative resistance, and consequently restoration of the infectivity or aggressiveness of pathogen population. The economic costs induced by pathogen adaptation may be huge, due firstly to the yield losses directly associated with an epidemic, and secondly because of the significant research and breeding efforts required to identify new resistance sources and develop new resistant cultivars [12]. Several deployment strategies have been proposed to improve the cost-effectiveness of plant resistance and prevent the frequently documented breakdown of major genes after their uniform deployment over large areas [3, 4, 9, 13, 14]. These strategies rely on the use of quantitative resistance, which is thought to be more durable because it poses smaller selection pressure on pathogen populations [1], or on the management of host genetic diversity [15–17]. This diversity can be introduced in time through crop rotations (e.g. recurring succession of different crops in the same field [18]). In space, different crops can be combined in the same field in cultivar mixtures [19, 20] or in different fields of the landscape as mosaics [3, 17]. Finally, several resistance sources can be stacked in the same cultivar through pyramiding [21, 22].

Few empirical studies have directly compared the performance of these different strategies, probably owing to the difficulty of implementing landscape scale experiments in practice (but see [23] for a comparison of all these strategies, except mosaics, using plastic tunnels). Models are not constrained by this difficulty, but surprisingly there are currently no published models enabling a global comparison. Indeed, most models focus on a single strategy (e.g. crop rotation [24, 25]; mixtures [26–28]; pyramiding [29, 30]; mosaics [31–42]) or a combination of strategies (e.g. mixture and pyramiding: [43]; mosaic and pyramiding: [44, 45]). Only a few studies explicitly compare two types of strategies [46–50] and only two studies evaluated more than two strategies [51, 52]. As a result, a comprehensive evaluation of different deployment schemes is complicated, and currently only feasible via pairwise comparisons [53, 54]. The situation for quantitative resistance is similar, since often only one [28, 34, 41, 42, 55], two [36, 37, 56], or a combination [26, 44, 49] of pathogen aggressiveness components are targeted, although quantitative resistance can affect several life-history traits of the pathogen. As articulated above, this current gap in our ability to predict which strategy will maximise our ability to control disease epidemics as well as pathogen evolutionary potential (or indeed whether these goals are compatible) emphasises the need for models that can compare different deployment schemes within the same framework, using standardised assumptions.

Comparison of different resistance deployment strategies requires the use of relevant evaluation criteria, which may vary depending on stakeholder objectives, and thus have an impact on the optimal strategy [39, 45]. Most published models focused only on one criterion, like resistance durability (i.e. the duration from initial deployment to the time when resistance is considered to have been overcome) (e.g. [30, 37, 47, 51, 52]), or epidemiological protection (i.e. reduction in pathogen population size and consequently in the proportion of diseased plants) (e.g. [26, 28, 32, 34–36, 43, 49, 50]). However, no correlation seems to exist between durability and epidemiological protection [45, 57], and those objectives can even be incompatible in severe epidemic contexts [33]. It is therefore essential to develop methods to assess deployment strategies against multiple evolutionary and epidemiological criteria [3, 17, 58].

The present study describes a demo-genetic model (i.e. it includes pathogen population demographic dynamics and its genetic evolution) which simulates the spread of a pathogen in a spatially explicit agricultural landscape. The model is flexible and can simulate mosaics, mixtures, rotation, and pyramiding of different major genes and up to four traits for quantitative resistance, acting on different components of pathogen aggressiveness (infection rate, latent period, infectious period, reproduction rate). Performance of resistance deployment is evaluated using several criteria from evolutionary (e.g. durability) or epidemiological (e.g. disease severity) perspectives. Thus, the model enables direct comparison of a range of spatio-temporal deployment strategies, but also enables investigation of the effects of landscape, epidemiological and evolutionary parameters on the ability of a given strategy to control disease. Although the main purpose of this paper is to provide a comprehensive description of the simulation model, we take advantage of the generality of the model and address three specific questions of interest to the scientific community. These questions aim to assess the potential of the combination of qualitative and quantitative resistance, given that the former prevents disease spread but is prone to breakdown, whereas the latter allows some disease development but to a lesser extent:

1. What is the durability of a major gene alone or combined with another source of resistance?
2. What is the level and speed of erosion of quantitative resistance alone or combined with a major gene?
3. What is the severity of epidemics in the landscape when qualitative and quantitative resistances are deployed alone or in combination?

The model can be parameterised to simulate various pathogen life histories. Here, we investigate the above questions in the context of rust diseases of cereal crops, caused by fungi of the genus *Puccinia* which can dramatically affect crop yields worldwide [59, 60]. Over the past decades, breeders have been engaged in an arms race against these pathogens, which exhibit high evolutionary potential in terms of their ability to overcome resistant crop cultivars following deployment [45, 60–63].

## Model

### Overview of the model

The model is stochastic, spatially explicit (the basic spatial unit is an individual field), based on a SEIR (‘susceptible-exposed-infectious-removed’) structure with a discrete time step. It simulates the spread (through clonal reproduction and dispersal) and evolution (via mutation) of a pathogen in an agricultural landscape, across cropping seasons split by host harvests which represent potential bottlenecks to the pathogen. The model is based on the model described in previous articles [40, 45], but has been considerably modified and extended to enable simulation of a wide array of deployment strategies: mosaics, mixtures, rotations and pyramiding of multiple major resistance genes which affect pathogen infectivity, and up to four quantitative resistance traits. These traits target different aggressiveness components of the pathogen, i.e. the infection rate, the duration of the latent period and the infectious period, and the propagule production rate. Initially, the pathogen is not adapted to any source of resistance, and is only present on susceptible hosts. However, through mutation, it can evolve and may acquire infectivity genes (which leads to breakdown of major resistance genes) or increase aggressiveness (which leads to the erosion of the relevant quantitative resistance traits). Evolution of a pathogen toward infectivity or increased aggressiveness on a resistant host is often penalised by a fitness cost on susceptible hosts [2, 64–66]. Consequently, in the present model, pathogens carrying infectivity genes may have lower infection rates (cost of infectivity) on susceptible hosts relative to pathogens that do not carry these genes. Similarly, a gain in pathogen aggressiveness on quantitatively resistant hosts is penalised by decreased aggressiveness on susceptible hosts, leading to a trade-off.

The evolutionary outcome of a deployment strategy is assessed by measuring the time until the pathogen achieves the three steps to adapt to plant resistance: (d1) first appearance of adapted mutants, (d2) initial migration to resistant hosts and infection (also referred as ‘arrival’ or ‘introduction’ in invasion biology), and (d3) broader establishment in the resistant host population (i.e. the point at which extinction becomes unlikely). Epidemiological outcomes are evaluated using the Green Leaf Area (GLA) as a proxy for yield, and the area under the disease progress curve (AUDPC) to measure disease severity.

### Landscape and resistance deployment strategies

In this study, a cropping landscape is represented by both its physical structure (defined as the spatial arrangement of field boundaries) and its genetic composition (defined by the allocation of crop cultivars within and among individual fields).

#### Landscape structure

Both real and simulated landscape structures can be used as input to the model. In this study, the landscape structure is simulated using a T-tessellation algorithm [67] (see S1 Figure) in order to control specific features such as number, area and shape of the fields, as described in previous studies [40, 45].

#### Allocation of crop cultivars

We used an algorithm based on latent Gaussian fields to allocate two different crop cultivars across the simulated landscapes (e.g. a susceptible and a resistant cultivar, denoted as SC and RC, respectively). This algorithm allows control of the proportions of each cultivar in terms of surface coverage, and their level of spatial aggregation (). Briefly, a random vector of values is drawn from a multivariate normal distribution with expectation 0 and a variance-covariance matrix which depends on the pairwise distances between the centroids of the fields. Next, the crop cultivars are allocated to different fields depending on whether each value drawn from the multivariate normal distribution is above or below a threshold. The proportion of each cultivar in the landscape is controlled by the value of this threshold. The sequential use of this algorithm allows the allocation of more than two crop cultivars (see S2 Figure). Therefore, deployment strategies involving two sources of resistance can be simulated by: (1) running the allocation algorithm once to segregate the fields where the susceptible cultivar is grown, and (2) applying one of the following deployment strategies to the remaining candidate fields:

i. Mosaics: two resistant cultivars (RC_1_ and RC_2_, carrying the first and the second resistance sources, respectively) are assigned to candidate fields by re-running the allocation algorithm;
ii. Mixtures: both RC_1_ and RC_2_ are allocated to all candidate fields;
iii. Rotations: RC_1_ and RC_2_ are alternatively cultivated in candidate fields, depending on the number of cropping seasons over which a given cultivar is grown before being rotated;
iv. Pyramiding: all candidate fields are populated with RC_12_, a resistant cultivar carrying both resistance sources.

Using this approach, a wide range of strategies can be simulated, noting that RC_1_ and RC_2_ are not necessarily deployed in balanced proportions or at a specific level of aggregation (i.e. these can also be varied to explore their effects). The same approach can be used to deploy more than two resistance sources. Alternatively, any crop allocation model (e.g. [68]) could be used to provide a more realistic landscape composition.

### Host-pathogen genetic interaction

All model parameters and values used in the simulations are listed in Table 1 (see S1 Text for details on model parameterisation to rust pathogens).

**Table 1.** Summary of the parameters used in the model and values for rust pathogens.

| Notation | Parameter | Value(s) for rust pathogens used in the simulations <sup>a</sup> |
| --- | --- | --- |
| **Simulation parameters** |  |  |
| Y | Number of simulated years | 50 years |
| T | Number of time steps in a cropping season | 120 days.year<sup>-1</sup> |
| J | Number of fields in the landscape | {155; 154; 152; 153; 156} <sup>b</sup> |
| V | Number of host cultivars | 2 |
| **Initial conditions and seasonality** |  |  |
| $C_v^0$ | Plantation host density of cultivar v | 0.1 m<sup>-2</sup> <sup>c</sup> |
| $C_v^{max}$ | Maximal host density of cultivar v | 2 m<sup>-2</sup> <sup>c</sup> |
| $\delta_v$ | Host growth rate of cultivar v | 0.1 day<sup>-1</sup> <sup>c</sup> |
| $\phi$ | Initial probability of infection | 5.10<sup>-4</sup> |
| $\lambda$ | Off-season survival probability | 10<sup>-4</sup> |
| **Pathogen aggressiveness components** |  |  |
| $e_{max}$ | Maximal expected infection rate | 0.40 spore<sup>-1</sup> |
| $\gamma_{min}$ | Minimal expected latent period duration | 10 days |
| $\gamma_{var}$ | Variance of the latent period duration | 9 days |
| $\Upsilon_{max}$ | Maximal expected infectious period duration | 24 days |
| $\Upsilon_{var}$ | Variance of the infectious period duration | 105 days |
| $r_{max}$ | Maximal expected propagule production rate | 3.125 spores.day<sup>-1</sup> |
| **Pathogen dispersal** |  |  |
| $g(.)$ | Dispersal kernel | Power-law function <sup>d</sup> |
| a | Scale parameter | 40 |
| b | Width of the tail | 7 |
| $\pi(.)$ | Contamination function | Sigmoid curve |
| $\kappa$ | Related to position of the inflexion point | 5.33 <sup>e</sup> |
| $\sigma$ | Related to position of the inflexion point | 3 <sup>e</sup> |
| **Host-pathogen genetic interaction** |  |  |
| G | Total number of major genes | {1; 2} |
| $\tau_g$ | Mutation probability for infectivity gene g <sup>f</sup> | 10<sup>-4</sup> <sup>g</sup> |
| $\tau_w$ | Mutation probability for aggressiveness component w <sup>f</sup> | 10<sup>-4</sup> <sup>g</sup> |
| $Q_w$ | Number of pathotypes relative to aggressiveness component w | 6 <sup>g</sup> |
| $\rho_g$ | Efficiency of major gene g | 1.0 <sup>g</sup> |
| $\rho_w$ | Efficiency of quantitative trait w | 0.5 <sup>g</sup> |
| $\theta_g$ | Cost of infectivity of infective gene g | 0.5 <sup>g</sup> |
| $\theta_w$ | Cost of aggressiveness for component w | 0.5 <sup>g</sup> |
| $\beta_w$ | Trade-off strength for aggressiveness component w | 1.0 <sup>g</sup> |

<sup>a</sup> model parameterisation is detailed in S1 Text.
<sup>b</sup> values for the five landscape structures.
<sup>c</sup> same value for all cultivars (no cost of resistance).
<sup>d</sup> $g(\|z' - z\|) = \frac{(b-2)(b-1)}{2\pi \cdot a^2} \cdot \left(1 + \frac{\|z' - z\|}{a}\right)^{-b}$ with $\|z' - z\|$ the Euclidian distance between locations z and z' in fields i and i', respectively; the mean dispersal distance is given by: $\frac{2a}{(b-3)} = 20$ m.
<sup>e</sup> the position of the inflexion point of the sigmoid curve is given by the relation $x_0 = ((\sigma - 1)/\kappa\sigma)^{1/\sigma} \approx 0.5$
<sup>f</sup> probability for a propagule to change its infectivity or its aggressiveness on a resistant cultivar carrying major gene $g$ or quantitative resistance trait $w$ .
<sup>g</sup> same value for all major genes, quantitative resistance traits, infectivity genes and aggressiveness components.

#### Host genotype

A host genotype (indexed by v) is represented by a set of binary variables, indicating the resistances it carries, denoted as mg_g_ (g=1,…,G) for major genes and qr_w_ (w=e,*γ*,r,γ, see also below) for quantitative resistance traits against the main components of pathogen aggressiveness.

#### Pathogen genotype

A pathogen genotype (indexed by p) is represented by a set of binary variables indicating whether it carries infectivity genes (variables ig_g_) and four discrete variables giving the level of aggressiveness for each life-history trait (variables ag_w_). All of these infectivity genes and aggressiveness components are assumed to evolve independently from each other.

#### Infectivity matrix

The interaction between potential major host resistance genes and associated pathogen infectivity genes is represented by a multiplicative factor of the infection rate of the pathogen. Thus, for major gene g, the possible interactions are summarised in an infectivity matrix (denoted as INF_g_, Table 2).

**Table 2.** Infectivity matrix.

| Infectivity matrix for<br>major gene $g$<br>$INF_g$ | | Host genotype $v$ | |
| --- | --- | --- | --- |
| | | Susceptible (SC)<br>$mg_g(v)=0$ | Resistant (RC)<br>$mg_g(v)=1$ |
| Pathogen<br>genotype $p$ | Non-infective<br>$ig_g(p)=0$ | 1 | $1 - \rho_g$ |
| | Infective<br>$ig_g(p)=1$ | $1 - \theta_g$ | 1 |

In the infectivity matrix, ρ_g_ is the efficiency of major resistance gene g on a non-adapted pathogen (ρ_g_ = 1 for a typical qualitative resistance gene conferring immunity [69], but ρ_g_ < 1 if a major gene is only partially expressed [3]), and θ_g_ is the cost of infectivity paid by an adapted pathogen on a susceptible host (θ_g_=0 means absence of cost of infectivity, θ_g_=1 means loss of infectivity on the susceptible cultivar).

#### Aggressiveness matrix

As for the infectivity matrices, the four potential traits for quantitative resistance (in the host) and the associated components for aggressiveness (in the pathogen) lead to four similar aggressiveness matrices (denoted as AGGw for component w, Table 3). These matrices summarise the multiplicative effect of this interaction on the infection rate of the pathogen (w=e), the duration of the infectious period (w=γ) and the propagule production rate (w=r), or the divisor effect on the duration of the latent period (w=*γ)*.

**Table 3.** Aggressiveness matrix.

| Aggressiveness matrix<br>for component w<br>$AGG^w$ | | Host genotype v | |
| --- | --- | --- | --- |
| | | Susceptible (SC)<br>$qr_w(v)=0$ | Resistant (RC)<br>$qr_w(v)=1$ |
| Pathogen genotype p | Non-aggressive<br>$ag_w(p)=1$ | 1 | $1 - \rho_w$ |
|  | ... | ... | ... |
|  | ... | ... | ... |
| | Fully aggressive<br>$ag_w(p)=Q_w$ | $1 - \theta_w$ | 1 |

Here, ρ_w_ is the efficiency of quantitative resistance with respect to aggressiveness component w for a non-adapted pathogen, θ_w_ the aggressiveness cost paid by adapted pathogens on susceptible hosts, and Q_w_ the number of possible pathotypes (thus, quantitative resistance trait w is completely eroded in Q_w_-1 steps). For non-adapted pathogens, every gain in aggressiveness on resistant hosts is penalised by a cost on susceptible hosts, according to the relationship [70]:

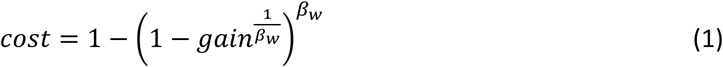

with β_w_ the strength of the trade-off for aggressiveness component w (Fig 2). β_w_<1 means a weak trade-off (gain higher than cost), β_w_>1 means a strong trade-off (gain lower than cost) and β_w_=1 represents a linear trade-off (gain equals cost). Thus, the aggressiveness associated with a pathogen genotype can be described by: (1 − *ρ*_*w*_) + *gain* × *ρ*_*w*_ on resistant hosts; and by: 1 − *cost* × *θ*_*w*_ on susceptible hosts.

**Fig 1.**
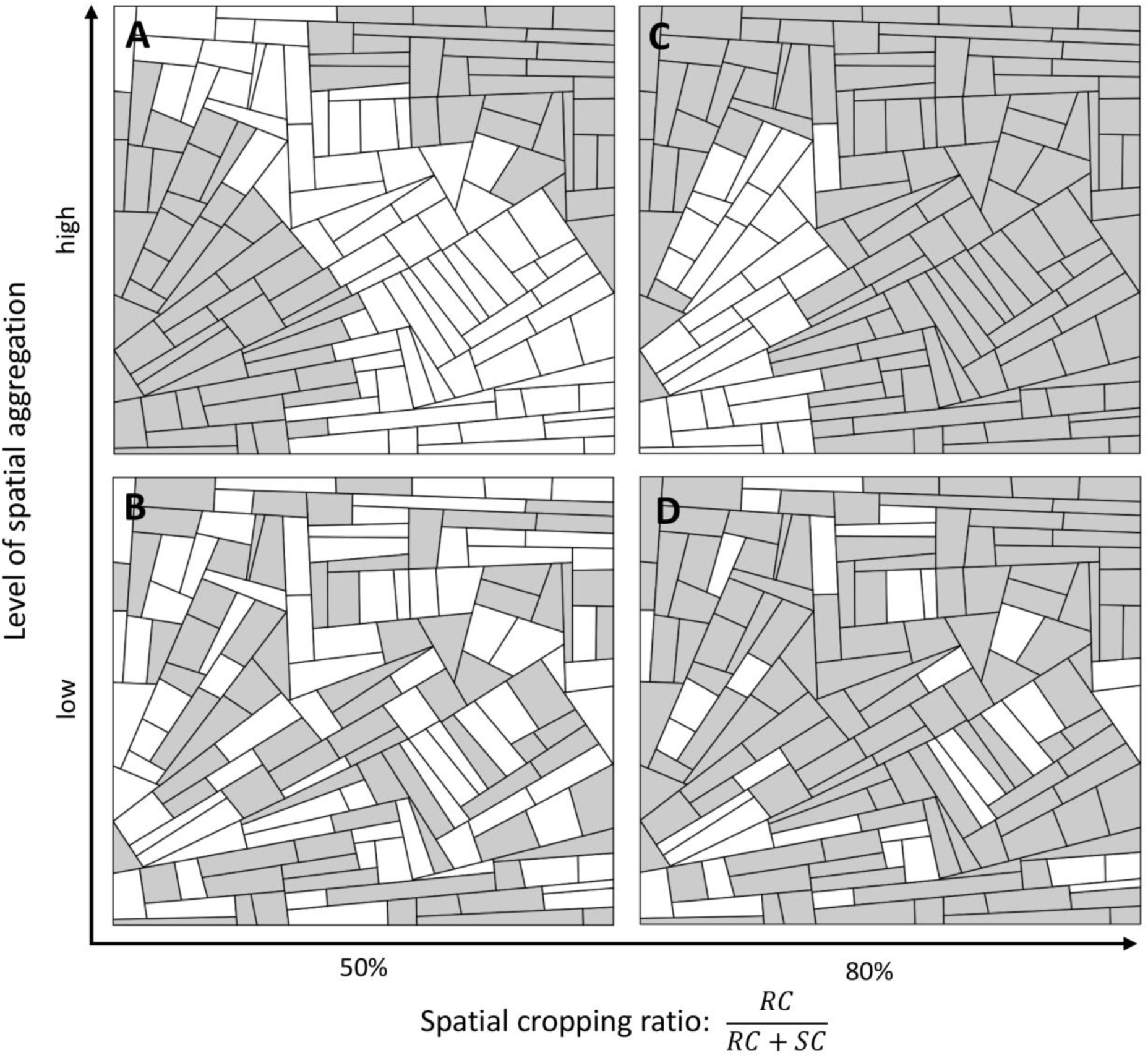
Allocation of two cultivars in the landscape. Firstly, a landscape structure is generated using T-tessellations. Secondly, the susceptible cultivar (SC, white) and resistant cultivar (RC, grey) are allocated to fields. Both the proportions of the surface coverage (horizontal axis: 50% in A and B, 80% in C and D) and level of spatial aggregation (vertical axis: high in A and C, low in B and D) for each cultivar are controlled via model input parameters.

**Fig 2.**
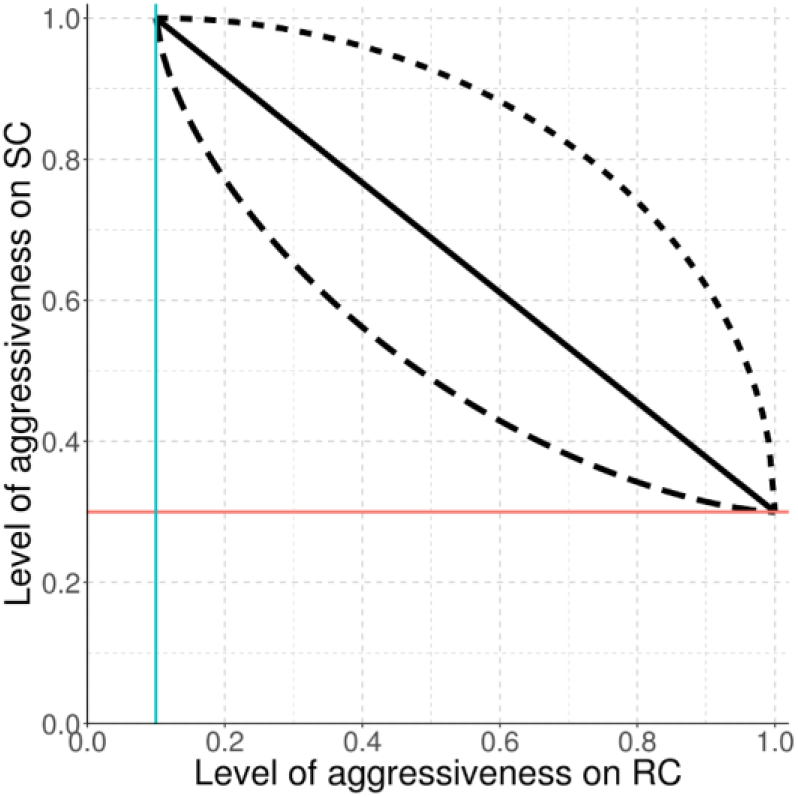
Trade-off relationship. Levels of pathogen aggressiveness on resistant (RC) and susceptible (SC) cultivars are linked by a linear (solid curve, β=1), a strong (dashed curve, β=1.5 in this example), or a weak (dotted curve, β=0.5) trade-off. The blue vertical line is related to resistance efficiency (ρ_w_=0.9) and the red horizontal line is related to the cost of aggressiveness (θ_w_=0.7).

Note, there are Q_w_ different pathotypes relative to aggressiveness component w, whereas there are only two pathotypes relative to each of the major infectivity genes. Thus the total number of pathogen genotypes is *P* = 2^*G*^ × ∏_*w*=*e,γ*,γ,*r*_ *Q*_*w*_. It is also noteworthy that the infectivity matrix (Table 2) shown here is a specific case of the more general aggressiveness matrix (Table 3 with Q_w_=2).

### Host and pathogen demo-genetic dynamics

The demo-genetic dynamics of the host-pathogen interaction are based on a SEIR structure. However, to avoid any confusion with the ‘susceptible’ cultivar, we have labelled this structure HLIR for ‘healthy-latent-infectious-removed’. Thus, in the following, H_i,v,t_, L_i,v,p,t_, I_i,v,p,t_, R_i,v,p,t_, and Pr_i,p,t_ respectively denote the number of healthy, latent, infectious and removed individuals (in this model, an ‘individual’ is a given amount of plant tissue, and is referred to as a ‘host’ hereafter for simplicity), and pathogen propagules, in field i (i=1,…,J), for cultivar v (v=1,…,V), pathogen genotype p (p=1,…,P) at time step t (t=1,…,TxY). T is the number of time steps in a cropping season and Y the number of simulated years (i.e. cropping seasons). Since the host is cultivated, we assume there is no host reproduction, dispersal or natural mortality (leaf senescence near the end of the cropping season is considered as part of host harvest). Fig 3 gives a schematic representation of the model structure.

**Fig 3.**
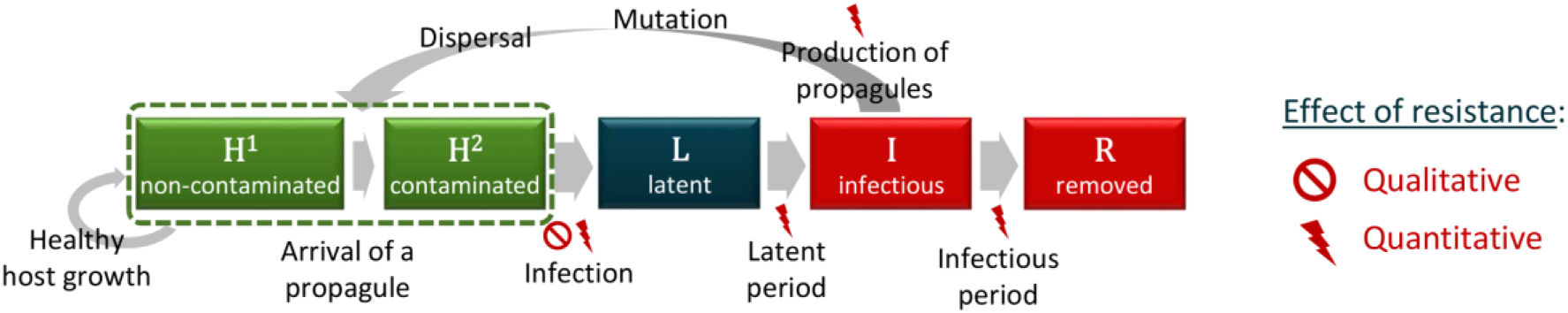
Architecture of the simulation model. Healthy hosts can be contaminated by propagules and may become infected. Following a latent period, infectious hosts start producing new propagules which may mutate and disperse across the landscape. At the end of the infectious period, infected hosts become epidemiologically inactive. Qualitative resistance usually prevents transition to the infected state, whereas quantitative resistance can affect several steps of the epidemic cycle but does not completely prevent infection. Green boxes indicate healthy hosts which contribute to crop yield and host growth, in contrast to diseased plants (i.e. symptomatic, red boxes) or those with latent infections (dark blue box).

#### Host growth

Only healthy hosts (denoted as *H*_*i,v,t*_) are assumed to contribute to growth of the crop (note, the model is flexible enough to relax this assumption; see Discussion). Thus, at each step t during a cropping season, the plant cover of cultivar v in field i increases as a logistic function, and the new amount of healthy plant tissue is:

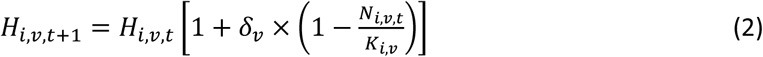

with δ_v_ the growth rate of cultivar v (possibly, although not necessarily, lower for resistant cultivars due to fitness costs of resistance [71]); 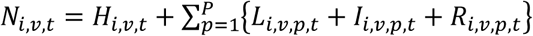 the total number of hosts in field i for cultivar v and at time t; and 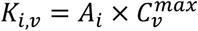 the carrying capacity of cultivar v in field i, which depends on A_i_, the area of the field, and 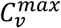, the maximal density for cultivar v. Note that when a mixture of several cultivars is present in the same field, decreased growth due to susceptible plants being diseased is not compensated by increased growth of resistant plants.

#### Contamination of healthy hosts

The healthy compartment (H) is composed of hosts which are free of pathogen propagules (H^1^), as well as hosts contaminated (but not yet infected) by the arrival of such propagules (H^2^). At the beginning of each step, all healthy hosts are considered free of propagules (H^1^). Then, as described in a previous modelling study [42], at time t in field i and for cultivar v, the number of contaminable hosts (i.e. accessible to pathogen propagules, denoted as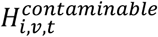) depends on the proportion of healthy hosts 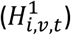 in the host population (*N*_*i,v,t*_):

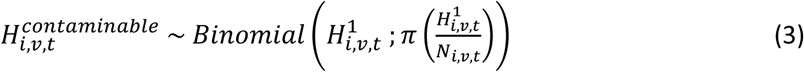

with 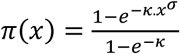, a sigmoid function with π(0)=0 and π(1)=1, giving the probability for a healthy host to be contaminated. Here, we assume that healthy hosts are not equally likely to be contacted by propagules, for instance because of plant architecture. Moreover, as the local severity of disease increases, eventually the probability for a single propagule to contaminate a healthy host declines due to the decreased availability of host tissue.

Following the arrival of propagules of pathogen genotype p in field i at time t (denoted as *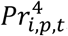*, see below), susceptible hosts become contaminated. The pathogen genotypes of these propagules are distributed among contaminable hosts according to their proportional representation in the total pool of propagules. Thus, for cultivar v, the vector describing the maximum number of contaminated hosts by each pathogen genotype (denoted as 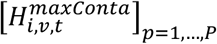 is given by a multinomial draw:

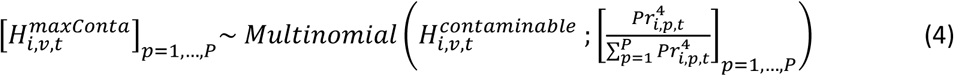

However, the number of deposited propagules 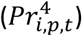 may be smaller than the maximal number of contaminated hosts 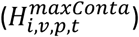. Thus, the true number of hosts of cultivar v, contaminated by pathogen genotype p in field i at t (denoted as 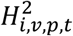) is given by:

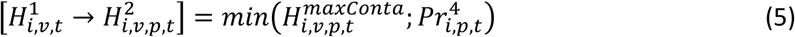

#### Infection

Between t and t+1, in field i, contaminated hosts 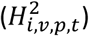 become infected (state L) with probability e_v,p_, which depends on the maximum expected infection efficiency, e_max_, and the interaction between host (v) and pathogen (p) genotypes:

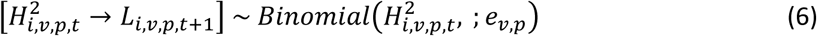

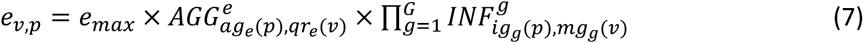

#### Latent period

Infected hosts become infectious (state I) after a latent period (LI) drawn from a Gamma distribution (a flexible continuous distribution from which durations in the interval [0; +∞[can be drawn) parameterised with an expected value, γ_exp v,p_, and variance, γ_var_. The expected duration of the latent period depends on the minimal expected duration, γ_min_, and the interaction between host (v) and pathogen (p) genotypes:

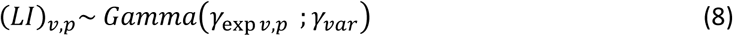

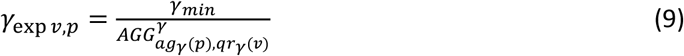

Note, the usual shape and scale parameters of a Gamma distribution, β_1_ and β_2_, can be calculated from the expectation and variance, exp and var, with: 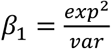 and 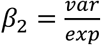, respectively.

#### Infectious period

Finally, infectious hosts become epidemiologically inactive (i.e. they no longer produce propagules, thus are in state R, ‘removed’) after an infectious period (IR) drawn from a Gamma distribution parameterised with expected value, γ_exp v,p_ and variance, γ_var_, similar to the latent period. The expected duration of the infectious period depends on the maximal expected duration, γ_max_, and the interaction between host (v) and pathogen (p) genotypes:

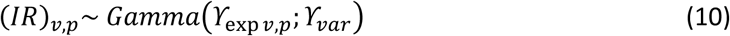

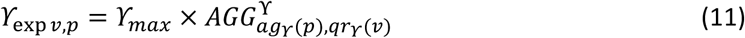

#### Pathogen reproduction

In field i at time t, infectious hosts associated with pathogen genotype p produce a total number of propagules (denoted as 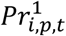), drawn from a Poisson distribution whose expectation, r_exp v,p_, depends on the maximal expected number of propagules produced by a single infectious host per time step, r_max_, and the interaction between host (v) and pathogen (p) genotypes:

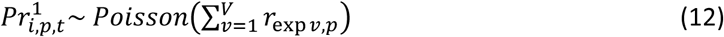

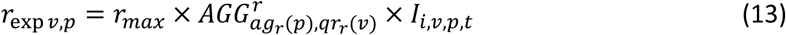

#### Pathogen mutation

The following algorithm is repeated independently for every potential infectivity gene g:

1. the pathotype (i.e. the level of adaptation with regard to major gene g, indexed by q; q=1,…,Q_g_; with Q_g_=2 since the pathotype is either infective, or non-infective) of the pathogen propagules is retrieved from their genotype p;
2. propagules can mutate from pathotype q to pathotype q’ with probability 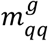, such as 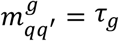 if q’≠q (hence 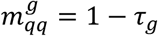 since Q_g_=2). Thus, in field i at time t, the vector of the number of propagules of each pathotype arising from pathotype q (denoted as 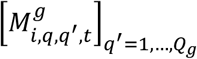) is given by a multinomial draw:

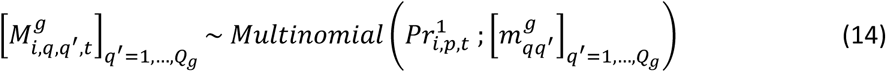
3. the total number of propagules belonging to pathotype q’ and produced in field i at time t (denoted as 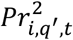) is:

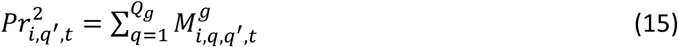
4. the new propagule genotype p’ is retrieved from its new pathotype (q’), and the number of propagules is incremented using a variable denoted as 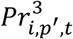.

The same algorithm is also repeated independently for every potential aggressiveness component w. In this case, τ_g_ is replaced by τ_w_ and Q_g_ by Q_w_. In addition, pathogen evolution is assumed to be stepwise, i.e. in a single step, a given pathotype can only mutate to a closely related pathotype (with regard to aggressiveness component w). Thus, we have 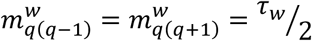 and 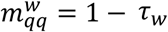 The exceptions are pathotypes fully adapted to either the susceptible host or the resistant host. These mutate towards less specialised pathotypes with a probability of τ_w_ to ensure their overall mutation probability is equivalent to that of other genotypes. In this model, it should be noted that the mutation probability τ_g_ (or τ_w_) is not the classic ‘mutation rate’ (i.e. the number of genetic mutations per generation per base pair), but the probability for a propagule to change its infectivity (or aggressiveness) on a resistant cultivar carrying major gene g (or quantitative resistance trait w). This probability depends on the classic mutation rate, the number and nature of the specific genetic mutations required to overcome major gene g (or improve aggressiveness component w), and the potential dependency between these mutations.

#### Pathogen dispersal

Propagules can migrate from field i (whose area is A_i_) to field i’ (whose area is A_i’_) with probability μ_ii’_, computed from:

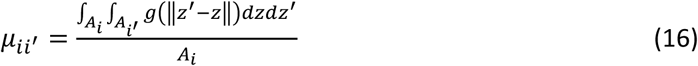

with ‖*z*, − *z*‖ the Euclidian distance between locations z and z’ in fields i and i’, respectively, and g(·) the two-dimensional dispersal kernel of the propagules. The computation of this probability is performed using the CaliFloPP algorithm [72]. Thus, at time t, the vector of the number of propagules of genotype p migrating from field i to each field i’ (denoted as 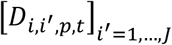 is:

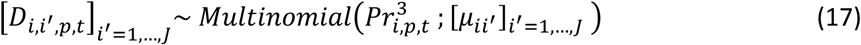

and the total number of propagules arriving in field i’ at time t (denoted as 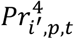 is:

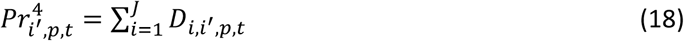

We consider that propagules landing outside the boundaries of the simulated landscape are lost (absorbing boundary condition), and there are no propagule sources external to the simulated landscape.

#### Seasonality

Let *t*^0^(*y*) and *t*^f^(*y*) denote the first and last days of cropping season y (y=1,…,Y), respectively. The plant cover in field i for cultivar v at the beginning of cropping season y is set at 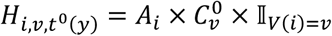, with 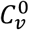 the plantation density of cultivar v and II_*V*_(_*i*_) an indicative variable set at 1 when field i is cultivated with cultivar v and 0 otherwise. At the end of a cropping season, the host is harvested. We assume that the pathogen needs a green bridge to survive the off-season. This green bridge could, for example, be a wild reservoir or volunteer plants remaining in the field (e.g. owing to incomplete harvest or seedlings). The size of this reservoir imposes a bottleneck for the pathogen population. The number of remaining infected hosts in field i for cultivar v and pathogen genotype p (denoted by 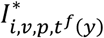 at the end of the off-season is given by:

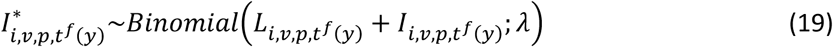

with λ the survival probability of infected hosts. Depending on host (v) and pathogen (p) genotypes, the number of propagules produced by the remaining hosts during their whole infectious period (denoted by 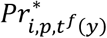) is drawn from a Poisson distribution:

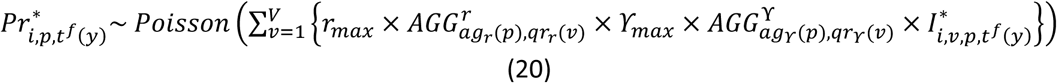

These propagules can mutate and disperse exactly as happens during the cropping season, and constitute the initial inoculum for the next cropping season.

#### Initial conditions

Atthe beginning of a simulation, healthy hosts are planted in each field as previously described. The initial pathogen population is assumed to be totally non-adapted to the resistance host cultivars, and is only present in susceptible fields with probability ϕ. Then the initial number of infectious hosts in these fields is: *I*_*i,v*=1,*p*=1,*t*=1_ = *Binomial*(*H*_*i,v*=1,*t*=1_; *ϕ*).

The model utilises multiple criteria to enable the assessment of different deployment strategies with regard to both evolutionary and epidemiological outcomes.

### Evolutionary outputs

#### Durability of qualitative resistance

For a given major gene, several computations are performed: (d1) time to first appearance of a pathogen mutant; (d2) time to first true infection of a resistant host by such mutants; and (d3) time when the number of infections of resistant hosts by these mutants reaches a threshold above which mutant pathogens are unlikely to go extinct. These metrics characterise the three critical steps to the breakdown of qualitative resistance: mutation toward infectivity, immigration (or introduction) on resistant hosts through dispersal, and subsequent broader establishment in the resistant host population.

#### Erosion of quantitative resistance

Since pathogen adaptation to quantitative resistance is gradual, the three measures described above are computed for every step towards complete erosion of resistance (i.e. Q_w_-1 levels). This allows us to characterise the point in time when quantitative resistance starts to erode, the final level of erosion, and the speed of erosion (i.e. the percentage of erosion per time unit from the time when quantitative resistance begins to erode).

#### Durability of a deployment strategy

A simulation run is divided into three periods: (1) the initial short-term period when all resistance sources are at their highest potential; (2) a transitory period during which a given deployment strategy is only partially effective; and (3) a longer-term period when all the resistances have been overcome or completely eroded. To assess the end of the short-term period, the time to establishment (durability measure (d3)) is computed for every major gene, and every quantitative trait at the first level of erosion (ag_w_(p)=2). The minimal value of these measures, denoted by D_1_, delimitates short-term and transitory periods. Similarly, the time to establishment is computed for every major gene, and for every quantitative trait at the highest level of erosion (ag_w_(p)=Q_w_). The maximal value of these measures, termed D_2_, delimits transitory and long-term periods. Fig 4 provides an example of a simulation run and the delimitation of these periods.

**Fig 4.**
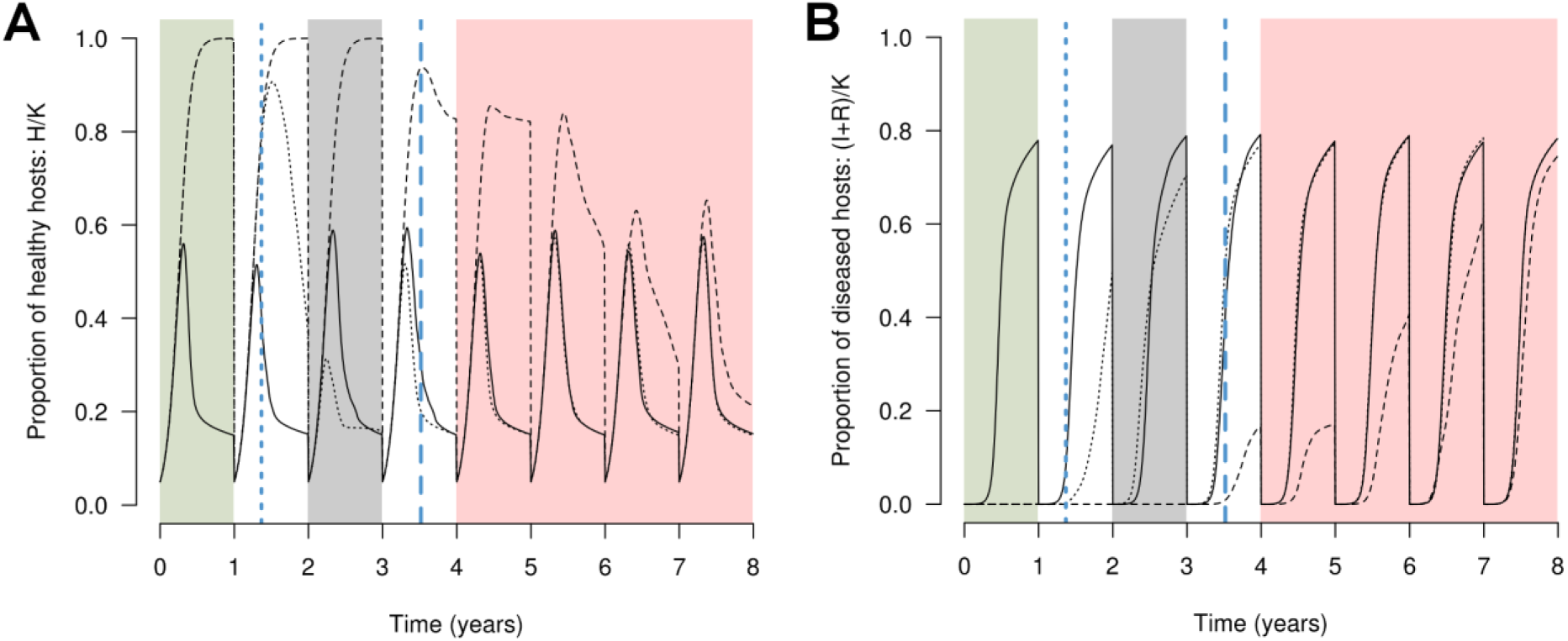
Computation of output variables in a simulated example. Two major resistance genes are deployed as a mosaic composed of a susceptible cultivar (solid curve) and two resistant cultivars (dotted and dashed curves) carrying the two major genes. The dynamics of the proportion of healthy (A) or diseased (B) hosts is integrated every year into the Green Leaf Area (GLA) or the area under disease progress curve (AUDPC), respectively. The vertical blue lines mark the times to breakdown of the first (dotted line) and the second (dashed line) major genes. These time points delimit the short-term (green zone), transitory (grey) and long-term (red) phases of resistance breakdown.

### Epidemiological outputs

The epidemiological impact of pathogen spread is evaluated by two different measures. Firstly, we use a measure termed Green Leaf Area (GLA), based on the Healthy Area Duration initially developed by Waggoner et al. [73]. The GLA represents the average number of productive hosts per time step and per surface unit, and is considered as a proxy for crop yield [74]. Secondly, we use the area under the disease progress curve (AUDPC), which is the average proportion of diseased hosts relative to the carrying capacity and represents disease severity [74]. We assume only heathy hosts (state H) contribute to crop production (Fig 3). Thus, for cultivar v during year y:

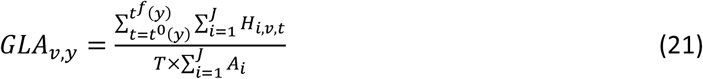

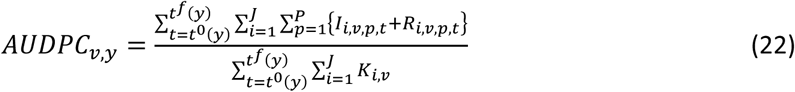

#### Global epidemiological control

The GLA and AUDPC of every cultivar (GLA_v_ and AUDPC_v_) as well as the whole landscape (GLA_TOT_ and AUDPC_TOT_) are averaged across the whole simulation run, to measure the global epidemiological performance of a deployment strategy.

#### Short-term epidemiological control

The average GLA and AUDPC of the susceptible cultivar is computed on whole cropping seasons from the beginning of the simulation until the end of the season preceding D_1_ (see ‘Evolutionary outputs’ for the computation of D_1_ and D_2_), and denoted as GLA_ST_ and AUDPC_ST_, respectively (Fig 4, green zone). These outputs cannot be computed if a resistance is overcome or starts to erode before the end of the first cropping season.

#### Epidemiological control during the transitory period

The average GLA and AUDPC of the susceptible cultivar is computed on whole seasons from the beginning of the season following D_1_ to the end of the season preceding D_2_, and denoted as GLA_TP_ and AUDPC_TP_, respectively (Fig 4, grey zone). These outputs cannot be computed if there is not at least one complete season between D_1_ and D_2_.

#### Long-term epidemiological control

The average GLA and AUDPC of the whole landscape is computed on whole seasons from the beginning of the season following D_2_ to the end of the simulation, and denoted as GLA_LT_ and AUDPC_LT_, respectively (Fig 4, red zone). These outputs cannot be computed if at least one of the resistances is not overcome or completely eroded by the end of the simulation.

### Application case: Durability and efficiency of combinations of different resistance sources

In this section, ten scenarios are simulated using the model to assess the evolutionary and epidemiological outcomes of deploying different pyramided combinations of qualitative and quantitative resistances. In this context, the model was parameterised to approximate biotrophic foliar fungal diseases as typified by rusts of cereal crops, caused by fungi of the genus *Puccinia*.

#### Model parameterisation for rust diseases

Within these pathosystems, propagules (called ‘spores’) are produced by sporulating lesions which develop on the leaves of infected hosts after a latent period of a few days, following which they are dispersed by wind. Thus, in this case, an ‘individual host’ in the model can be considered as a foliar site where a spore can land and potentially trigger the development of a localised infection. The parameter values associated with pathogen aggressiveness components (infection rate, latent period duration, sporulation duration, sporulation rate) were estimated using available data from the literature [29, 30, 33, 37, 39–42, 45, 50, 55, 56, 60, 65, 75–116]; other parameters were arbitrarily fixed (see Table 1 for parameter values and S1 Text for details). In addition, the time step was set at one day and simulations were run over 50 cropping seasons of 120 days per year. Given this parameterisation, the threshold for mutant pathogen establishment (and thus resistance breakdown) was set at 50,000 infections. Above this threshold, the probability of extinction of a pathogen genotype is below 1% (see S2 Text for details of how the threshold was calculated).

#### Simulated scenarios

A set of five landscape structures of about 150 fields within a total area of 2×2 km^2^ was generated (see S1 Figure). Areas of the simulated fields ranged from 0.36 to 5.38 ha (mean: 2.60 ha). Of these fields, 80% were cultivated with a resistant cultivar, and the remaining 20% with a susceptible cultivar, with a low level of spatial aggregation (landscape typified by Fig 1D). In the first five simulated scenarios, the resistant cultivar carried a single resistance source: either a major gene (conferring complete immunity against non-infective pathogens, ρ_g_=1, as typically described in gene-for-gene scenarios for many rusts [62]), or one of the four traits for quantitative resistance (against infection rate, latent period duration, sporulation rate, or sporulation duration). The efficiency of quantitative resistance traits was set at ρ_w_=0.5, meaning that infection rate, sporulation rate or the duration of the sporulation period of non-aggressive pathogens on resistant hosts was reduced by 50%. Similarly, the latent period duration was increased in such a way that the number of epidemic cycles of a non-aggressive pathogen in a cropping season was reduced by 50% on resistant hosts. In the next five scenarios, the previous resistance sources (a major gene or one of the four quantitative resistance traits) were combined with a major gene to represent a single pyramided cultivar. In all scenarios, the mutation probabilities were set at τ_g_=τ_w_=10^-4^ in such way that a cultivar carrying a single major gene would be overcome in less than one year. Each scenario was simulated ten times on all five landscape structures, resulting in 50 replicates per scenario, and a total of 500 simulations.

The model is written using the C and R languages, and is available via the R package landsepi (v0.0.2, [117]). Within the R software (v3.4.0, [118]) landscape structures were generated using the package RLiTe (http://kien-kieu.github.io/lite/rlite.html). The cultivar allocation algorithm used positive definite matrices generated with the *Exponential* function of the package fields (v8.10, [119]) and the *nearPD* function of the package Matrix (v1.2-11, [120]). The CaliFloPP algorithm was performed using the package RCALI (v0.2-18, [72]). Depending on the number of resistance sources, one simulation run takes 60 to 180 seconds on a standard desktop computer (2 cores, 2.30 GHz).

## Results

Several different scenarios were simulated to investigate three specific questions relative to the evolutionary and epidemiological outcomes of deploying combinations of qualitative and quantitative resistances against rust diseases of cereal crops. In this context, we varied landscape structure (i.e. spatial arrangement of field boundaries, see S1 Figure) but not its composition (i.e. proportion and spatial aggregation of the resistant cultivar). Landscape structure was never significant, thus in what follows we consider that every scenario was replicated 50 times.

### What is the durability of a major gene alone or combined with another source of resistance?

To assess the durability of a single major gene, the time period from the beginning of the simulation to appearance of mutants able to overcome the major gene (d1), first infection of the resistant cultivar (d2), and broader establishment on the resistant cultivar (d3) were used. Under the simulated conditions, mutants always appeared, dispersed across the landscape and established on a resistant cultivar carrying one major gene in less than one year (Fig 5A, ‘MG1’). When the resistant cultivar carried a second major gene (i.e. as a pyramid with two major genes), the first infection of a resistant host was delayed to 8.3 years (90% central range, CR_90_: 0.4-23.4) on average, and the pathogen population was not established before, on average, 20.7 years (CR_90_: 0.9-50.0) (Fig 5A, ‘MG2’).

**Fig 5.**
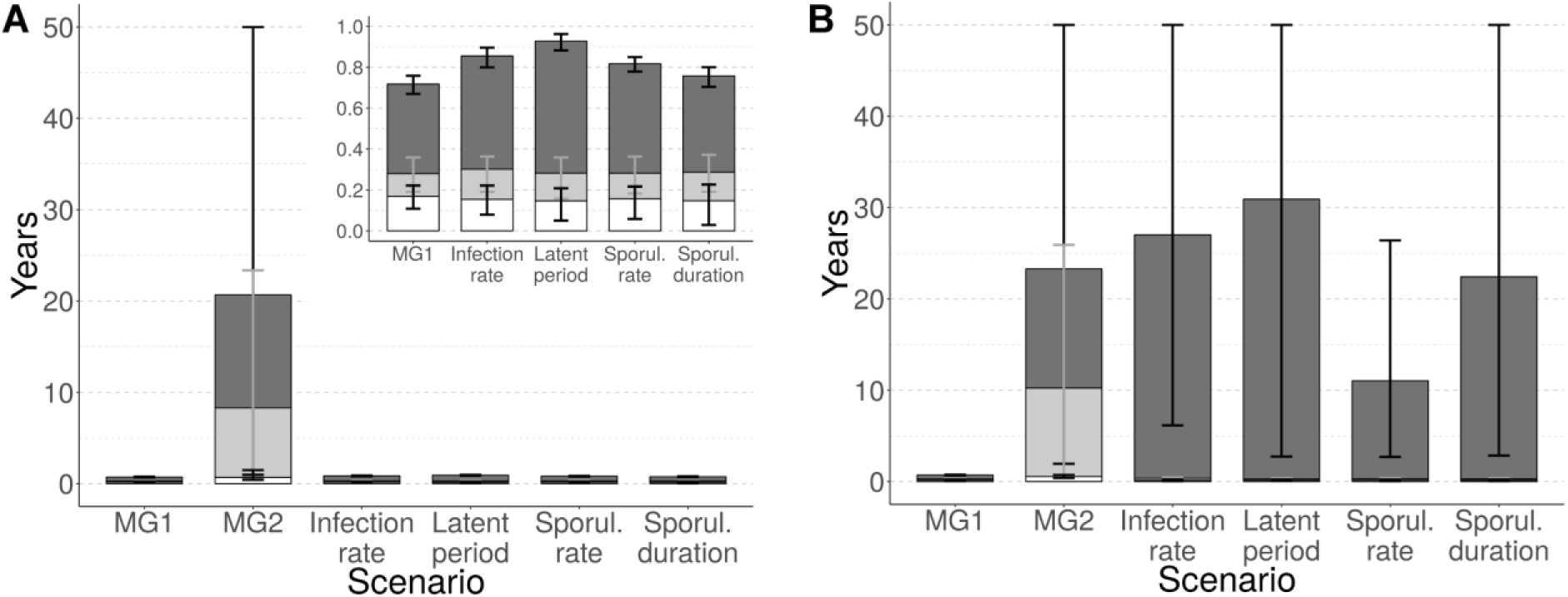
Evolutionary outcomes for *Puccinia* rusts after 50 years: durability of the major gene. Time to appearance of mutants carrying the associated infectivity gene (white segment of bars), to first infection (light grey) or to establish (dark grey) on resistant hosts. The resistant cultivar carries a single major gene (MG1, efficiency ρ_1_=100%), a combination of two major genes (MG2, efficiency ρ_2_=100%) or the single major gene is combined with one of several quantitative resistance traits (columns 3 to 6, efficiency ρ_w_=50% in A and ρ_w_=90% in B). Inset: enlargement of scenarios showing short durability. Every scenario is replicated 50 times. Vertical bars represent the 90% central range (represented in grey for the time to first infection, for legibility).

In contrast, when a major gene was combined with a quantitative resistance with a 50% efficiency, the breakdown of the major gene was nearly as quick as if the major gene was alone, regardless of the pathogen life-history trait targeted by the quantitative resistance (Fig 5A, column 3 to 6, and inset). This conclusion is consistent with the results of an experimental study on pepper resistance against root-knot nematode [121], but differs from those of other studies carried out on different pathosystems, showing that a quantitative resistant background can significantly increase the durability of a cultivar carrying a major gene for resistance [122–124]. In order to test if this difference could be due to our assumption that quantitative resistance has a 50% efficiency, we replicated the numerical experiment with higher efficiencies (ρ_w_=60%, 70%, 80%, 90%). The results of these new simulations indicated that when quantitative resistance efficiency is higher than 80%, durability of the major gene is improved in combination with quantitative resistance, especially if latent period is targeted (time to establishment delayed to 5.2 years, CR_90_: 1.7-10.8, see S3A Figure). Furthermore, above 90% efficiency, quantitative resistance can increase the durability of the major gene compared to pyramiding of two major genes (average time to establishment between 11.0 and 30.9 years depending on the targeted trait, Fig 5B).

### What is the level and speed of erosion of quantitative resistance alone or combined with a major gene?

When deployed alone, quantitative resistance was on average eroded by 26.4% (CR_90_: 20-40), 20.8% (CR_90_: 0-40), 22.0% (CR_90_: 0-40), and 19.2% (CR_90_: 0-40) by the end of the simulation, for resistances targeting respectively infection rate, duration of the latent period, sporulation rate and duration of the sporulation period (Fig 6). The associated average speeds of erosion were 2.16, 0.78, 1.44 and 1.02% per year, respectively. The targeted pathogen life-history trait appeared to have a significant effect on the final level (Kruskal Wallis χ^2^ tests with 3 degrees of freedom, p=8.10^-9^) and speed (p=6.10^-5^) of resistance erosion. The combination with a major gene significantly affected the final level (Kruskal Wallis χ^2^ test with 1 df, p=8.10^-3^), but not the speed (p=0.85) of erosion.

**Fig 6.**
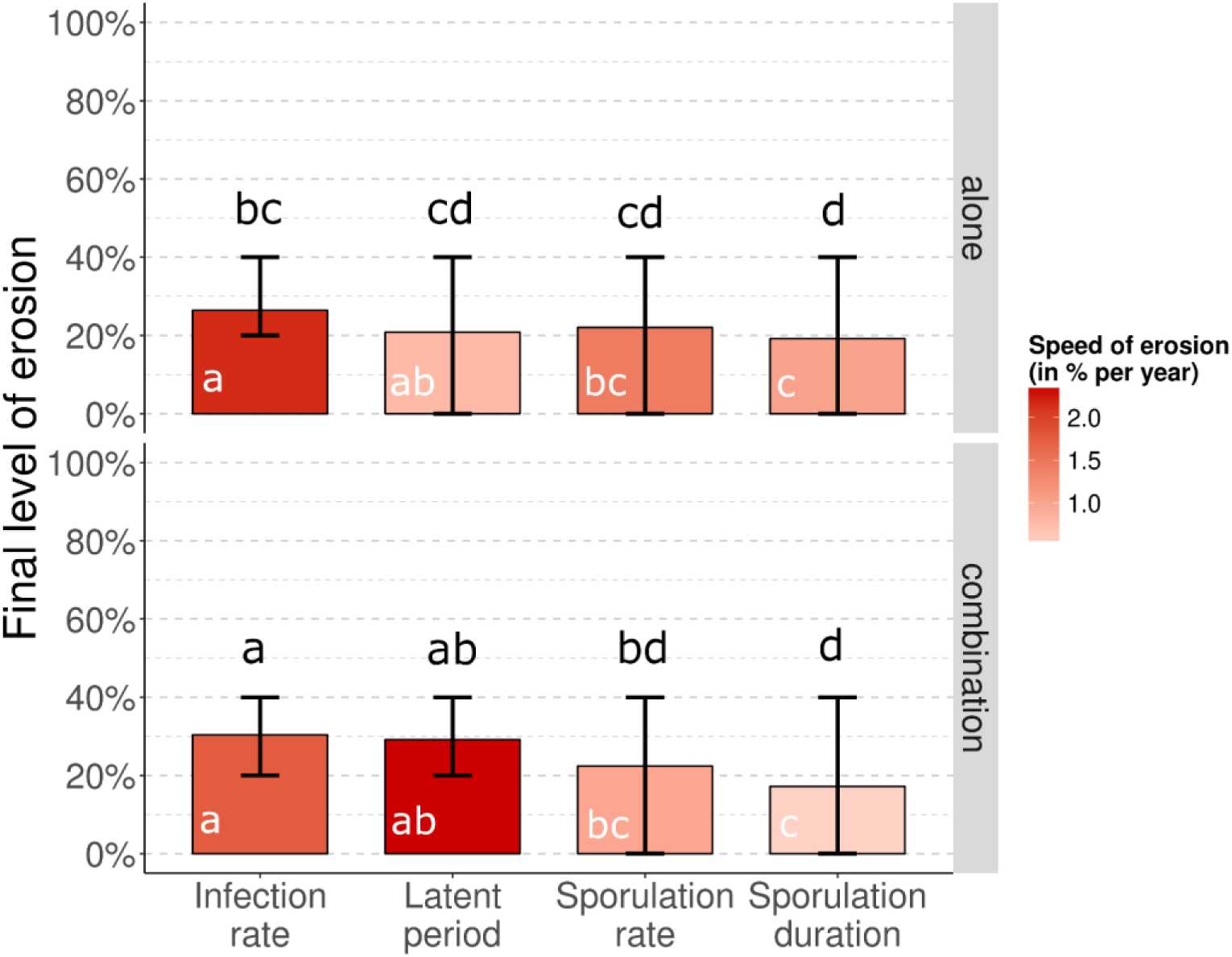
Evolutionary outcomes for *Puccinia* rusts after 50 years: Final level of erosion of quantitative resistance traits. Quantitative resistance (efficiency ρ_w_=50%) is deployed alone (top row) or in combination with a major resistance gene (bottom row). The red shading indicates the average speed of erosion from the time when quantitative erosion starts to erode to the time when the final level of erosion is reached, with darker shades representing faster erosion rates. Lower case letters indicate statistically different groups for final level (black) and speed (white) of erosion, according to pairwise comparison tests (Dunn’s test with Holm correction). Every scenario is replicated 50 times. Vertical bars represent the 90% central range.

It is important to remember that the speed of erosion was computed from the time point when quantitative resistance started to erode (hence, following breakdown of the major gene) and not from the beginning of simulations. Assessing the average speed of erosion from the beginning of the simulations would greatly impact the results if the major gene was durable for several years. As previously described, this is not the case with a 50% efficiency of quantitative resistance. However, with a 90% efficiency, the average speed of erosion of quantitative resistance from the beginning of the simulation was 2.5 to 3.0 % per year when deployed alone, and only 0.7 to 1.7 % per year when combined with a major gene. In this context, the combination of quantitative resistance with a major gene significantly delayed the start of quantitative resistance erosion (Kruskal Wallis χ^2^ test with 1 df, p<10^-15^).

### What is the severity of epidemics when qualitative and quantitative resistances are deployed alone or in combination?

The AUDPC of the susceptible cultivar (AUDPC_SC_), the resistant cultivar (AUDPC_RC_) and across the entire cropping landscape (AUDPC_TOT_) averaged across the whole simulation period were used as indicators of the severity of epidemics in the simulated scenarios (see S4 Figure for examples). As expected, regardless of the cultivar or the deployment scenario, disease severity in a landscape where a resistant cultivar is deployed was lower than in a fully susceptible landscape (dashed line in Fig 7).

**Fig 7.**
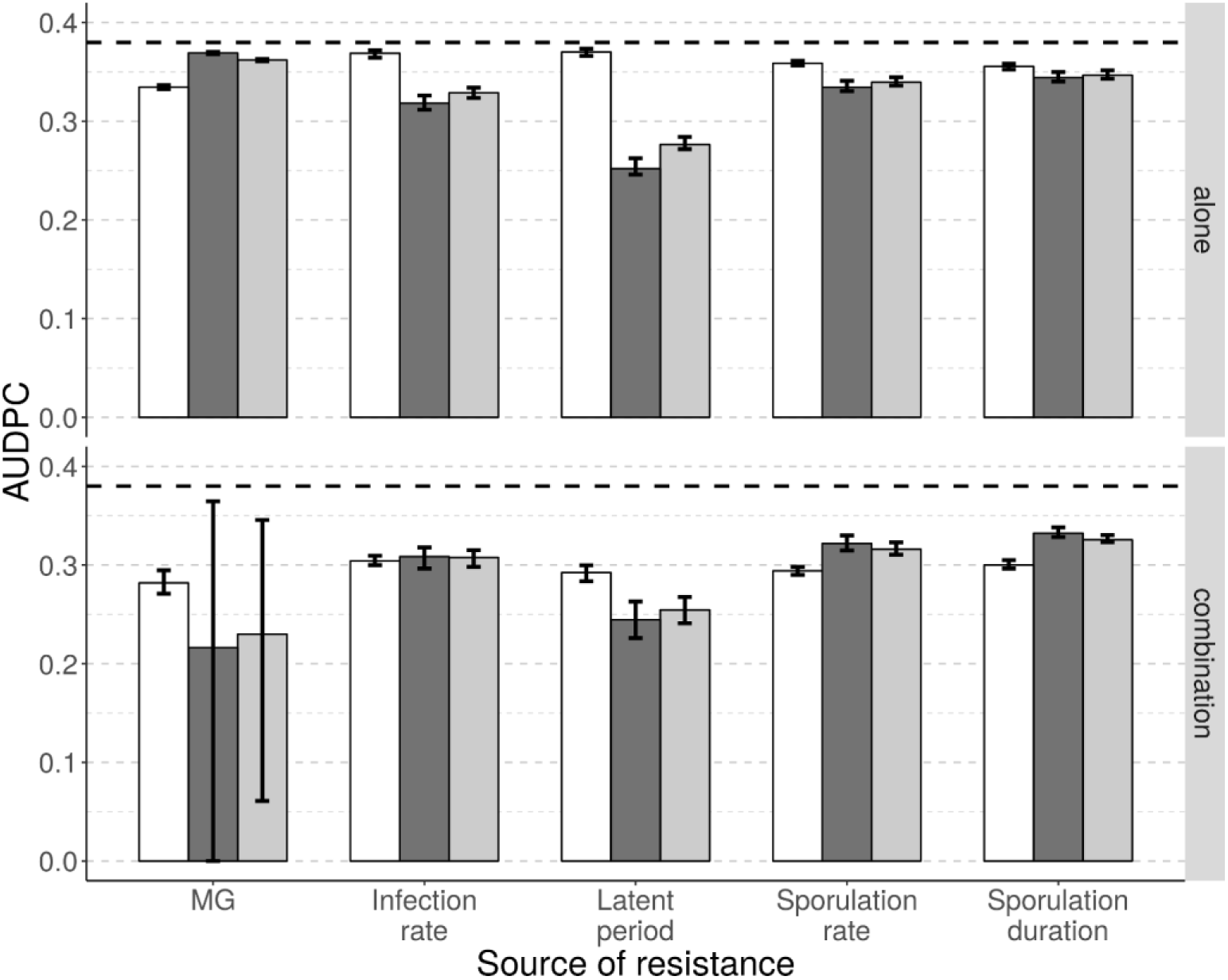
Epidemiological outcomes for *Puccinia* rusts after 50 years. Average area under disease progress curve (AUDPC) for a susceptible (white bars) and a resistant (dark grey bars, cropping ratio: 80%) cultivar, as well as for the whole landscape (light grey bars). The top row represents situations where the resistant cultivar carries a single major gene (MG, efficiency ρ_1_=100%) or a single quantitative resistance trait (columns 2 to 5, efficiency ρ_w_=50%). The bottom row shows results when a major resistance gene was added to each of the single resistance scenarios (e.g. the leftmost set of bars on the bottom row shows results for two major genes in combination; the rightmost set is when a major gene was combined with quantitative resistance against sporulation duration). Each scenario was replicated 50 times. The horizontal dashed line represent the average AUDPC in a fully susceptible landscape. Vertical bars represent the 90% central range.

When only a single resistance source was incorporated into the resistant cultivar, the most effective source was quantitative resistance against duration of the latent period (AUDPC_RC_=0.25), followed by quantitative resistance against infection rate, sporulation rate and duration of the sporulation period (AUDPC_RC_=0.32, 0.33 and 0.34, respectively), and finally major gene resistance (AUDPC_RC_=0.37). When a major gene was combined with one of these resistance sources, the combination of two major genes became the most efficient strategy (AUDPC_RC_=0.22) but also the most variable (CR_90_: 0.00-0.37). Below this, were combinations including quantitative resistance traits in the same order as above and with similar efficiencies (AUDPC_RC_=0.24, 0.31, 0.32, and 0.33 against latent period, infection rate, sporulation rate and sporulation duration, respectively).

For the susceptible cultivar, disease severity was less variable between scenarios than for the resistant cultivar. The average AUDPC_SC_ was 0.33, 0.37, 0.37, 0.36 and 0.36, respectively when a major gene or a quantitative resistance trait against infection rate, latent period, sporulation rate or duration of the sporulation period was deployed alone. However, when a major gene was combined with these sources, AUDPC_SC_ was smaller (0.28, 0.30, 0.29, 0.29 and 0.30, respectively).

Globally, across the whole landscape, disease severity (AUDPC_TOT_), was very similar to levels of disease severity seen in the resistant cultivar (AUDPC_RC_), since this cultivar constituted a high proportion of the area being cropped (80%, see Fig 1D for an example). The global severity of disease could also be assessed using the GLA (focusing on healthy hosts) instead of the AUDPC (focusing on diseased hosts). In our context, GLA_TOT_ was highly negatively correlated with AUDPC_TOT_ (Pearson correlation coefficient of −0.92, p<10^-15^).

## Discussion

This article describes the first spatiotemporal model with the ability to flexibly simulate the evolution of a plant pathogen in a cropping landscape, following the deployment of resistance within the four main categories of deployment strategies: mosaics, mixtures, rotations and pyramiding. In addition, the model includes every possible combination of different major genes (qualitative resistance conferring immunity to the plant) and traits for quantitative resistance against the main aggressiveness components of the pathogen (infection rate, latent period, infectious period, and reproduction rate). Because of its flexible parameterisation and ability to simulate an explicit landscape, the model offers the possibility to vary the number and type of deployed resistance sources, their relative proportion in surface coverage across the landscape and their level of spatial (or temporal) aggregation, as well as the epidemiology and the evolutionary potential of the pathogen. In addition, although the present study used a simulated landscape in order to control particular features of the fields, the model also permits the use of real landscapes to match more specific applied contexts. This model is implemented in the package *landsepi* [117] for the statistical computing environment R. Consequently, it provides a new and useful tool to assess the performance of a wide range of deployment options. In particular, it was used here to evaluate the combination of different sources of resistance, and it will be used in future studies to explore different spatio-temporal deployment strategies.

With respect to criteria used to evaluate the performance of different deployment strategies, these may vary depending on needs of different stakeholder groups (e.g. breeders and growers). Thus, the model generates a panel of outputs describing the epidemiological (i.e. the ability of different resistance sources to reduce the severity of epidemics and consequently their impact on crops) and the evolutionary (relative to the durability of these resistance sources) outcomes of a given deployment strategy.

Owing to similar epidemiological and evolutionary concepts, there are interesting parallels between the deployment of plant resistance to pathogens and the application of pesticides (for plants) or vaccines (for animals, including humans). Indeed, the proportion of resistant fields in our study is equivalent to the proportion of fields treated with a pesticide [29] or the proportion of vaccinated individuals in an animal population [125]. The efficiency of plant resistance is similar to pesticide dose [46], and the type of resistance may be compared to the type of pesticide or vaccine. For instance, quantitative resistance traits against pathogen infection rate, latent period and sporulation rate are analogous to imperfect vaccines with anti-infection, anti-growth or anti-transmission modes of action, respectively [126]. It is noteworthy that spatio-temporal deployment strategies for plant resistance have counterparts in the application of pesticides. Thus, mosaics of different resistant crops are analogous to mosaics of fields treated with different pesticides, rotations of resistant crops to periodic application of different pesticides, and crop mixtures and pyramids of resistance genes to combinations of pesticide molecules [53, 54].

The model is flexible enough to facilitate investigation of a wide range of host-pathogen systems. Although our application case focused on biotrophic rust fungi of cereal crops, other pathogens transmitted by wind, rain or vector insects (e.g. many fungi, bacteria and viruses), can be simulated by modifying in particular the dispersal kernel and the value of parameters associated with pathogen aggressiveness components (see for example how to parameterise the model to necrotrophic fungal pathogens in S1 Text). Nevertheless, some fixed features of the model (e.g. clonal reproduction of the pathogen) may need to be modified if sexual reproduction (or recombination for viruses) significantly contribute to pathogen genetic diversity and dynamics. In addition, modelling vector-borne bacteria and viruses transmitted in a persistent manner may require incorporation of vector dynamics and behaviour into the model framework. Future work on the model is planned to address some of these current limitations.

### A spatially explicit stochastic model of pathogen evolution

Three main properties distinguish models simulating the deployment of resistance and the interpretation of their results: whether they are demographic or demo-genetic, whether they are deterministic or stochastic, and whether they are spatial or not.

In contrast with purely demographic models, our model includes the genetic evolution of the pathogen in addition to pathogen population dynamics and host growth during a cropping season. This enables the explicit simulation of the appearance of new genotypes through mutation [29, 30, 46, 48], which represents the first step towards resistance breakdown or erosion (i.e. prior to migration to and broader establishment onto resistant hosts). The time required for the achievement of this first step can be the main determinant of the durability of some deployment strategies, like pyramiding [51]. This step is thus essential to comprehensively assess the relative performance of different deployment strategies.

We further note that the present model is stochastic, i.e. it relies on probabilistic computations of simulated biological processes. It is consequently well able to account for biologically realistic random events [127]. As illustrated by Lo Iacono et al. [35], who used a stochastic version of the model developed by van den Bosch et al. [39], the likelihood of extinction events in pathogen populations can considerably impact the performance of a deployment strategy.

Finally, because our model is able to simulate explicit landscapes, it accounts for spatial heterogeneity, which affects landscape connectivity and consequently the ability of the pathogen to disperse. Pathogen dispersal, in addition to being one of the unavoidable steps of adaptation to plant resistance, strongly shapes pathogen evolutionary dynamics [12, 41, 128]. Moreover, the spatial nature of this model enables a wide range of deployment options to be considered at different spatial scales, particularly given the possibility to vary deployment parameters like the proportion of the landscape across which resistant cultivars are planted [32, 51], as well as the level of spatial aggregation [28, 45, 48, 49, 129, 130].

### Computation of epidemiological and evolutionary outputs

Resistance durability (i.e. the period of time from initial deployment of a resistant cultivar to when resistance is considered to have been overcome) is a typical evolutionary target of resistance deployment strategies, but its computation in a model is not obvious. Proposed methods include targeting the point in time when the first adapted pathogens appear [29, 30, 39], when their prevalence [37, 48, 55, 131] or frequency in pathogen population [39, 51, 52] exceeds a threshold, or when productivity of the resistant cultivar drops below an arbitrary threshold [45].

Since different measures give different information, the present model includes several measures of durability: the time until first appearance of mutants, the time until first infection of a resistant host by such mutant, and the time until the prevalence of these mutants exceeds a threshold. The first measure assesses the ability of deployment strategies to reduce the probability of appearance of mutants by reducing pathogen population size. The difference between the first and the second measure provides information on the potential to hinder pathogen migration to resistant hosts, and the difference between the second and the last measure is related to the rate of establishment on resistant cultivars as a result of the balance between selection and genetic drift (occurring between seasons in this model). With respect to the establishment process, it is interesting to note that different stages of invasion can be targeted by changing the value of the threshold. Importantly, we considered that time to establishment was best computed using a prevalence threshold (i.e. total number of resistant hosts infected by adapted pathogens) as opposed to using a frequency-dependent threshold in the pathogen population (i.e. the proportion of adapted pathogens in the global pathogen population). This is because the frequency of adapted pathogens may never (or always) exceed a threshold simply as a result of a very low (or very high) proportion of resistant hosts in the landscape relative to susceptible hosts.

As pointed out in the introduction, evolutionary and epidemiological outcomes are not necessarily correlated. In this context, several epidemiological outputs are used in this study to characterise the level of protection provided by a deployment strategy against the potential damage caused by an epidemic. Some previous studies focused on the final state of the simulations or on the point at which a stable evolutionary equilibrium is reached, thus proposing criteria related to the final proportion of healthy [43, 44] or infected [26, 49, 129] hosts. However, in addition to long-term measures related to stable equilibria, short-term and transitory periods measures are important because severe epidemics responsible for heavy losses may occur during these stages [125, 132]. These periods can be accounted for by averaging the number of healthy hosts over the whole simulation run using an analogy of our Green Leaf Area [35–37, 40, 42, 133] or the number of infected hosts using the area under disease progress curve (AUDPC) [28, 32, 33, 48, 50]. Interestingly, these measures offer the possibility to concentrate on different evolutionary phases, for example the short-term period following resistance deployment until resistance breakdown, and the long-term period once resistance is overcome [45].

In the present study, the proposed outputs are based on both GLA and AUDPC, computed not only for the whole simulation run to have a global snapshot of the epidemiological outcome, but also for different time periods: from initial resistance deployment until the first resistance is overcome, from the time when all resistances are overcome until the end of the simulation as well as during the transitory period. The performance of various deployment strategies can differ during these three periods, since they are associated with different epidemiological contexts. Thus it is important to consider these different measures together. Moreover, the objectives of different stakeholders define whether such criteria should be computed from the AUDPC or from the GLA. Indeed, if the objective is to limit the amount of disease, the use of AUDPC-based variables may be more appropriate, since they target infected hosts. In contrast, GLA-based variables represent the amount of healthy hosts which generally represent the largest contribution to crop yield. Thus, this variable can be useful when considering the impact of a given deployment strategy on productivity, especially when resistance costs, different planting densities, or different host species are involved. It is interesting to note that here, only healthy hosts were assumed to participate in host growth and contribute to final yield, considering that the infection by a pathogen consumes host resources. For rust pathogens of cereal crops, this assumption seems reasonable since it has been shown that manual defoliation of wheat leaves decreases yield less than infection by *Puccinia striiformis* [73]. Nevertheless, this assumption can easily be changed by including hosts at different sanitary stages (e.g. latently infected but not yet diseased) in the logistic equation of host growth and in computation of the GLA.

### Combining different sources of resistances

In this study, we present an initial exploration of the model where we evaluate the potential of combining qualitative and quantitative resistance to control rust of cereal crops. We note that in this exploration some parameters (in particular mutation probabilities, see S1 Text) were arbitrarily fixed to study a simple and theoretical situation where a resistant cultivar carrying a single major gene would be rapidly overcome following deployment. This is because our intent in this case was to compare different resistance combinations rather than provide an absolute prediction of the durability and efficiency of a particular strategy. Regardless, altering the mutation probabilities, as long as they are the same for all infectivity genes and aggressiveness components, changes our results quantitatively but not qualitatively (see S1 Text and the results of simulation performed with smaller mutation probabilities in S5 Figure).

#### Durability of qualitative resistance

We found greater durability of a major resistance gene when combined with another major gene than when combined with a quantitative resistance trait exhibiting moderate efficiency. The high durability of pyramided major genes is often explained by the low probability of the pathogen simultaneously acquiring the required mutations to infect the resistant cultivar, added to the severe fitness costs associated with these mutations [64]. In the scenario we evaluated, it seems that this probability was not low enough to prevent the rapid appearance of adapted mutants (Fig 5A, ‘MG2’, durability measure (d1)). On the other hand, the great difference between durability measures (d1) and (d2) indicates that a long time passed before these mutants were able to disperse to resistant fields and infect a resistant host. This delay can be explained by the low probability that a spore simultaneously mutates and disperses to a resistant field, added to the severe fitness costs imposed on mutant pathogens in susceptible fields (note, in our simulations the infection rate of a pathogen carrying two infectivity genes is reduced by 75% on susceptible hosts). Furthermore, the great difference between durability measures (d2) and (d3) suggests that many extinction events impeded mutant establishment on the resistant cultivar, likely due to bottlenecks between cropping seasons.

Conversely, the combination of a major resistance gene with a highly efficient source of quantitative resistance can considerably increase the durability of the major gene. It is noteworthy that the durability of the major gene becomes also more variable (compare the 90% central ranges in Fig 5 and S3A Figure). This is mainly attributed to an increasing proportion of simulations where the major gene has a very long durability or is still not overcome by the end of the 50-year simulated period (e.g. in 42% of the simulations performed with a major gene combined with a quantitative resistance targeting pathogen latent period; see Fig 5B ‘Latent period’).

#### Durability of quantitative resistance

Our results show that the presence of a major gene in plant genotype does not greatly affect pathogen adaptation to a moderately effective quantitative resistance. However, highly effective quantitative resistance delays the breakdown of the major gene which at the same time delays the start of quantitative resistance erosion. Once quantitative resistance begins to erode, one would expect complete erosion over time, as observed in experimental serial passages performed with a plant virus [134]. In our results, quantitative resistance was not completely eroded by the end of the simulations. This was not due to the number of simulated years, since a speed of erosion of about 2% per year should enable complete erosion within 50 years of simulation. Moreover, it is interesting to note that quantitative resistance was eroded by approximately 25% when the initial resistance efficiency was 50%, and eroded by about 50% when the initial efficiency was 90% (Fig 6 and S3C Figure). This suggests that, in our simulations, pathogen adaptation always converged towards the same level of aggressiveness, which is likely a fitness optimum, given the fitness costs associated with pathogen adaptation, the shape of the trade-off function, and the proportion of the resistant cultivar in the landscape.

#### Epidemiological outcomes for combined qualitative and quantitative resistances

Complementing a moderately efficient quantitative resistance with a major gene seems to have a greater effect on disease severity on the susceptible cultivar than on the resistant cultivar. This is probably because under these conditions the major gene has a very short durability (see above and Fig 5A), and only slightly delays the erosion of quantitative resistance (see above and Fig 6). On the other hand, pathogen evolution towards increased infectivity and aggressiveness on the resistant cultivar (which is strongly favoured in our simulations since 80% of the landscape is grown with a resistant cultivar) triggers a stronger fitness cost on the susceptible cultivar, than for situations when only the evolution of increased aggressiveness is necessary to adapt to host resistance. This results in decreased severity on the susceptible cultivar and consequently across the whole landscape. Our results also suggest that quantitative resistance against the duration of the latent period has the best potential to limit disease severity, especially when the resistance is highly efficient (see S6B Figure). This is in agreement with previous studies [8, 93, 135], which highlighted the key role of the number of epidemic cycles within a cropping season.

#### General conclusions

As expected, the combination of multiple resistance sources in a resistant cultivar results in improved evolutionary and epidemiological outcomes relative to the deployment of a single resistance. In addition to the greater evolutionary barrier this represents, such combinations are more likely to trigger high fitness costs for the pathogen, thus constraining its ability to be simultaneously well adapted to different cultivars. Our simulations indicate that the pyramiding of two major genes is highly durable and effective, because this completely blocks the infection of resistant hosts by non-adapted pathogens, and requires the simultaneous acquisition of two costly infectivity genes to be overcome. Nevertheless, despite its partial efficiency, quantitative resistance may provide good epidemiological control if it has a sufficiently high efficiency (see S6 Figure) and especially if it is complemented by a major gene, which can significantly delay the time when quantitative resistance starts to erode. In turn, quantitative resistance helps reduce disease severity once the major gene is overcome.

Using a demographic model, Pietravalle et al. [37] found that quantitative resistance against the infection rate or the propagule production rate of the pathogen are equally efficient. However, in a demo-genetic version of a similar model, Lo Iacono et al. [36] found a greater effect of quantitative resistance against pathogen infection rate. Our results are consistent with this second study. However, our model includes for the first time the four main pathogen aggressiveness components. According to our simulation results, the most promising trait for quantitative resistance is against the duration of the latent period, owing to its influence on the number of epidemic cycles the pathogen can complete during a growing season. Consequently, quantitative resistance targeting pathogen latent period could effectively supplement the use of major genes in breeding programs. Thus, latent periods of rust pathogens could be measured on different host cultivars, and quantitative trait loci (QTL) associated with longer latent periods could be identified through genome-wide association studies, and subsequently selected for in marker-assisted selection programs [136].

It is important to remember that, based on our model assumptions and the values chosen for parameters associated with pathogen evolution, the scenario we simulated here was a relatively simple one. For example, we considered that the different quantitative traits evolve independently from each other by assuming that: (i) mutations are independent from each other; (ii) when a quantitative resistance trait is deployed, only the targeted aggressiveness component of the pathogen can evolve; and (iii) a fitness cost is paid on the susceptible cultivar on the same aggressiveness component. Nevertheless, there is considerable evidence that different pathogen aggressiveness components often vary in association with each other [8, 83], suggesting that they do not necessarily evolve independently. Several trade-offs between aggressiveness components have been described [6, 7], and such trade-offs may considerably impact the performance of a particular deployment strategy [56]. In addition, some aggressiveness component(s) not directly targeted by host resistance could counterbalance the affected component(s), like a compensation phenomenon (as simulated in some modelling studies [137, 138]), and the fitness cost on the susceptible cultivar could be paid on an aggressiveness component different from the evolving component. These aspects could be captured through the use of statistical estimates of the genetic variance and covariance of pathogen life-history traits, issued from phenotype values (G-matrix [139]). In addition, even in the case of well documented pathosystems like rusts of cereal crops, calibration of some parameters can be challenging (e.g. effective sporulation rate and dispersal kernel, see S1 Text), either because data obtained in laboratory experiments may not be representative of what happens in the field, or simply because data are missing. To fill this gap and help calibrate simulation models, estimation models built with exactly the same parameters can be useful to estimate poorly known parameters from field data [140–142].

### Future research directions

In a recent review dealing with the combination of qualitative and quantitative resistances, Pilet-Nayel et al. [143] wrote “There is still a need to adequately choose resistance QTLs to create optimal combinations and limit QTL erosion”. The present work highlights that longer latent periods may be one promising target. Nevertheless, the general features of the model could be used to test if our conclusions hold in different contexts. Our simulations showed that the efficiency of quantitative resistance (ρ_w_) has a strong impact on the durability and epidemiological control of plant resistance. Therefore, in future studies we plan to assess the influence of the efficiency of qualitative resistance by including partially effective major genes (ρ_g_<1, [3]). We also envisage varying the mutation probabilities (τ_g_ and τ_w_), cost of infectivity (θ_g_), cost of aggressiveness (θ_w_), and number of steps to erode a trait for quantitative resistance (Q_w_), to simulate various possible choices of major genes and traits for quantitative resistance. It would also be of interest to simulate and investigate the potential for combinations of several sources of quantitative resistance, whose performance on disease severity has been experimentally demonstrated [144]. Finally, we will also explore different spatiotemporal strategies to deploy plant resistance to pathogens. In particular, we plan to compare mosaics, mixtures, rotations and pyramids of genetic resistances with respect to their epidemiological and evolutionary outcomes. The challenge is to identify strategies for a given host-pathogen interaction that are not only durable and efficient, but also feasible and likely to be adopted. In this context, promising strategies identified with the simulation model could be experimentally tested in the field. In addition, broader consideration of the economic context will also help maximise the chances that identified strategies are considered for real deployment.

## Acknowledgements

The authors thank Loïc Houde for computing assistance, Jeremy Burdon for stimulating discussions and for reviewing this manuscript, as well as three anonymous reviewers for their constructive and helpful comments. Simulations were performed using the Pearcey cluster within the CSIRO supercomputer platform. L.R., L.G.B. and P.H.T. were funded by grant CSP00192 from the Grain Research & Development Corporation (GRDC, https://grdc.com.au/). All authors were supported by the INRA/CSIRO linkage program.

## Supporting information

**S1 Figure**. Landscape structures used in this study.

**S2 Figure**. Spatial allocation of three cultivars across a cropping landscape.

**S3 Figure**. Evolutionary outcome when quantitative resistance is highly effective.

**S4 Figure**. Dynamics of diseased hosts (AUDPC) from different scenarios of resistance deployment.

**S5 Figure**. Evolutionary and epidemiological outputs when the mutation probabilities are smaller.

**S6 Figure**. Epidemiological outcome when quantitative resistance is highly effective.

**S1 Text**. Model parameterisation to rust diseases.

**S1 Table**. Available data on the duration of latent and sporulation periods for rust diseases caused by fungi of the genus *Puccinia*.

**S2 Table**. Mean dispersal distance of rust spores computed from available results obtained in studies which used exponential functions.

**S7 Figure**. Distribution of the latent period duration and length of the sporulation period of rust diseases caused by fungi of the genus *Puccinia*.

**S8 Figure**. Two-dimensional representation of the power-law dispersal kernel used in this study.

**S9 Figure**. Sigmoid contamination function.

**S10 Figure**. Deterministic dynamics of healthy hosts in the absence of disease.

**S11 Figure**. Dynamics of healthy and diseased hosts in a fully susceptible landscape in a simulated example.

**S2 Text**. Calculation of the threshold for pathogen establishment.

**S12 Figure**. Probability of extinction of a mutant pathogen depending on the number of infections.

