## Supplementary Materials for "Assessing the durability and efficiency of landscape-based strategies to deploy plant resistance to pathogens"

The following supporting information is available for this article:

**S1 Figure.** Landscape structures used in this study.

**S2 Figure.** Spatial allocation of three cultivars across a cropping landscape.

**S3 Figure.** Evolutionary outcome when quantitative resistance is highly effective.

**S4 Figure.** Dynamics of diseased hosts (AUDPC) from different scenarios of resistance deployment.

**S5 Figure.** Evolutionary and epidemiological outputs when the mutation probabilities are smaller.

**S6 Figure.** Epidemiological outcome when quantitative resistance is highly effective.

**S1 Text.** Model parameterisation to rust diseases.

**S1 Table.** Available data on the duration of latent and sporulation periods for rust diseases caused by fungi of the genus *Puccinia.*

**S2 Table.** Mean dispersal distance of rust spores computed from available results obtained in studies which used exponential functions.

**S7 Figure.** Distribution of the latent period duration and length of the sporulation period of rust diseases caused by fungi of the genus *Puccinia.*

**S8 Figure.** Two-dimensional representation of the power-law dispersal kernel used in this study.

**S9 Figure.** Sigmoid contamination function.

**S10 Figure.** Deterministic dynamics of healthy hosts in the absence of disease.

**S11 Figure.** Dynamics of healthy and diseased hosts in a fully susceptible landscape in a simulated example.

**S2 Text.** Calculation of the threshold for pathogen establishment.

**S12 Figure.** Probability of extinction of a mutant pathogen depending on the number of infections.

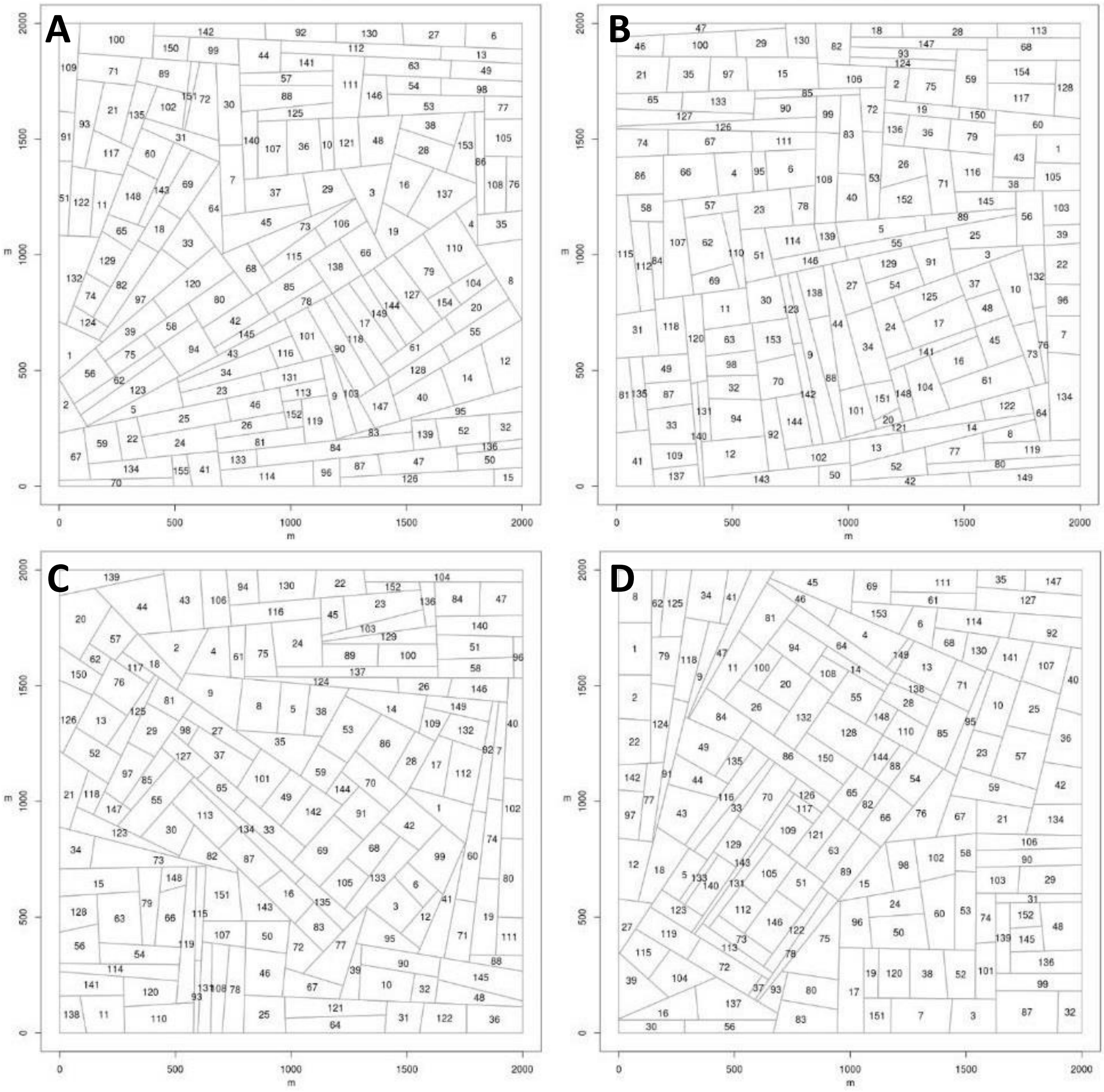


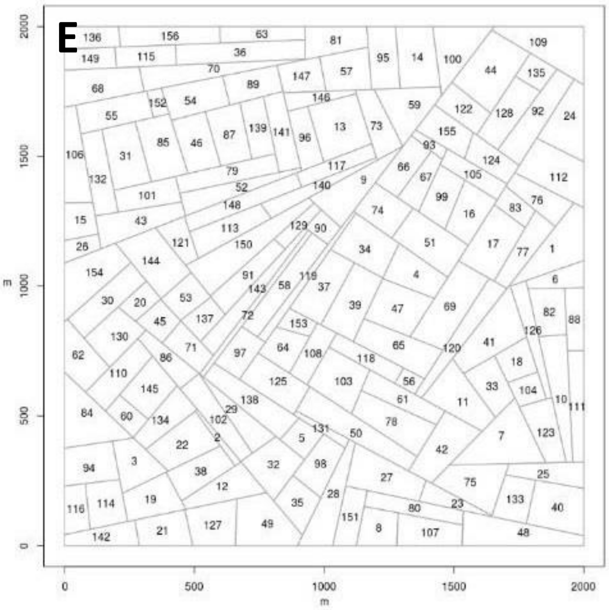


**S1 Figure. Landscape structures used in this study.** Landscapes of about 150 fields (total area: 2x2 km$^2$) were simulated using a T-tessellation algorithm from package RLiTe (http://kien-kieu.github.io/lite/rlite.html), with controlled number, area and shape of the fields. Numbers correspond to field indices. The effect of landscape structure was never statistically significant in our simulations.

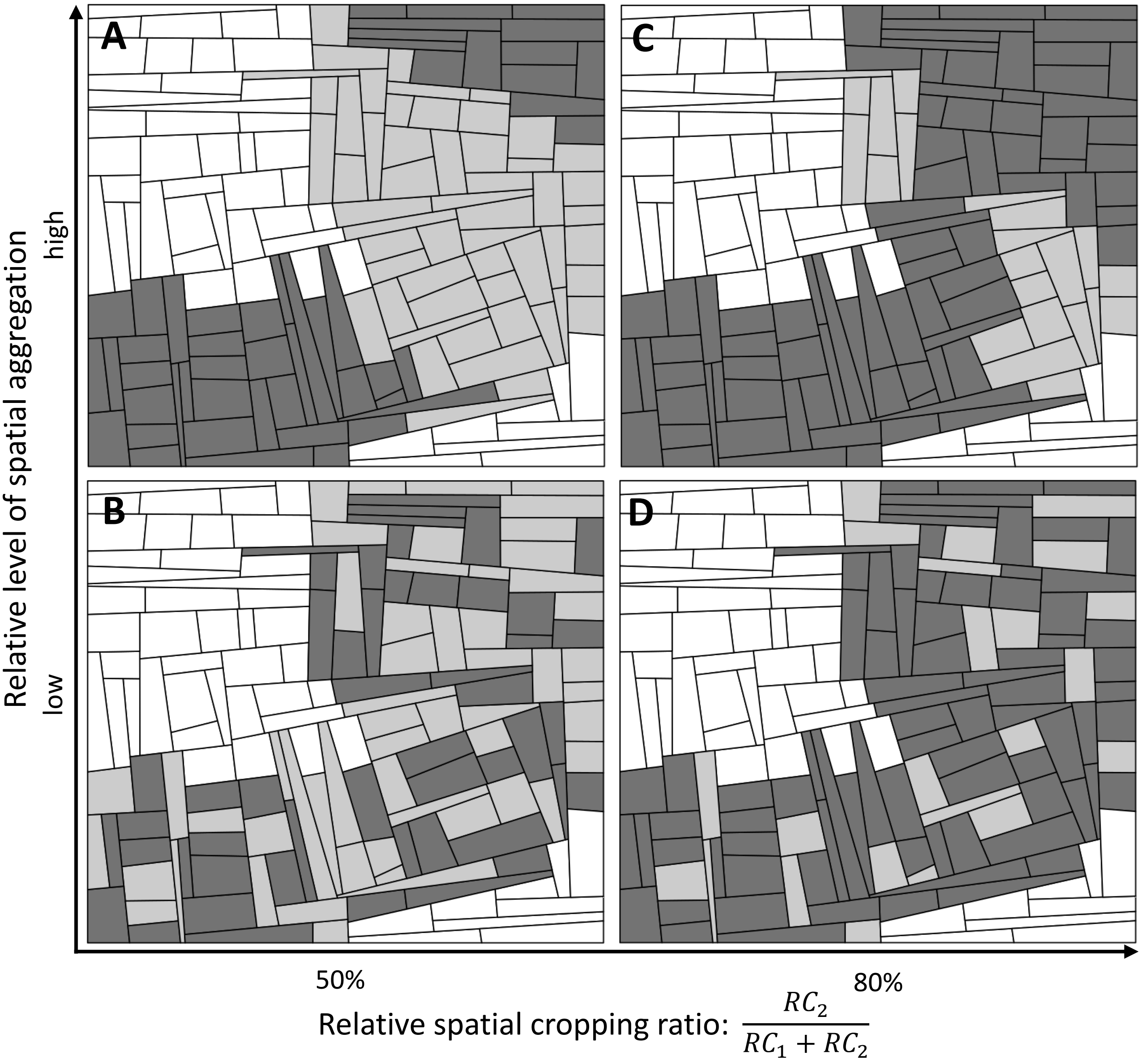


**S2 Figure. Spatial allocation of three cultivars across a cropping landscape.** A landscape structure is generated using T-tessellations. Then, a susceptible (white) and two resistant cultivars ($RC_1$, light grey, and $RC_2$, dark grey) are allocated to fields, with controlled relative proportions of the surface coverage (horizontal axis: 50% in A and B, 80% in C and D) and level of spatial aggregation (vertical axis: high in A and C, low in B and D) of $RC_1$ and $RC_2$. The total proportion of resistant fields is 2/3 and their level of aggregation is high.

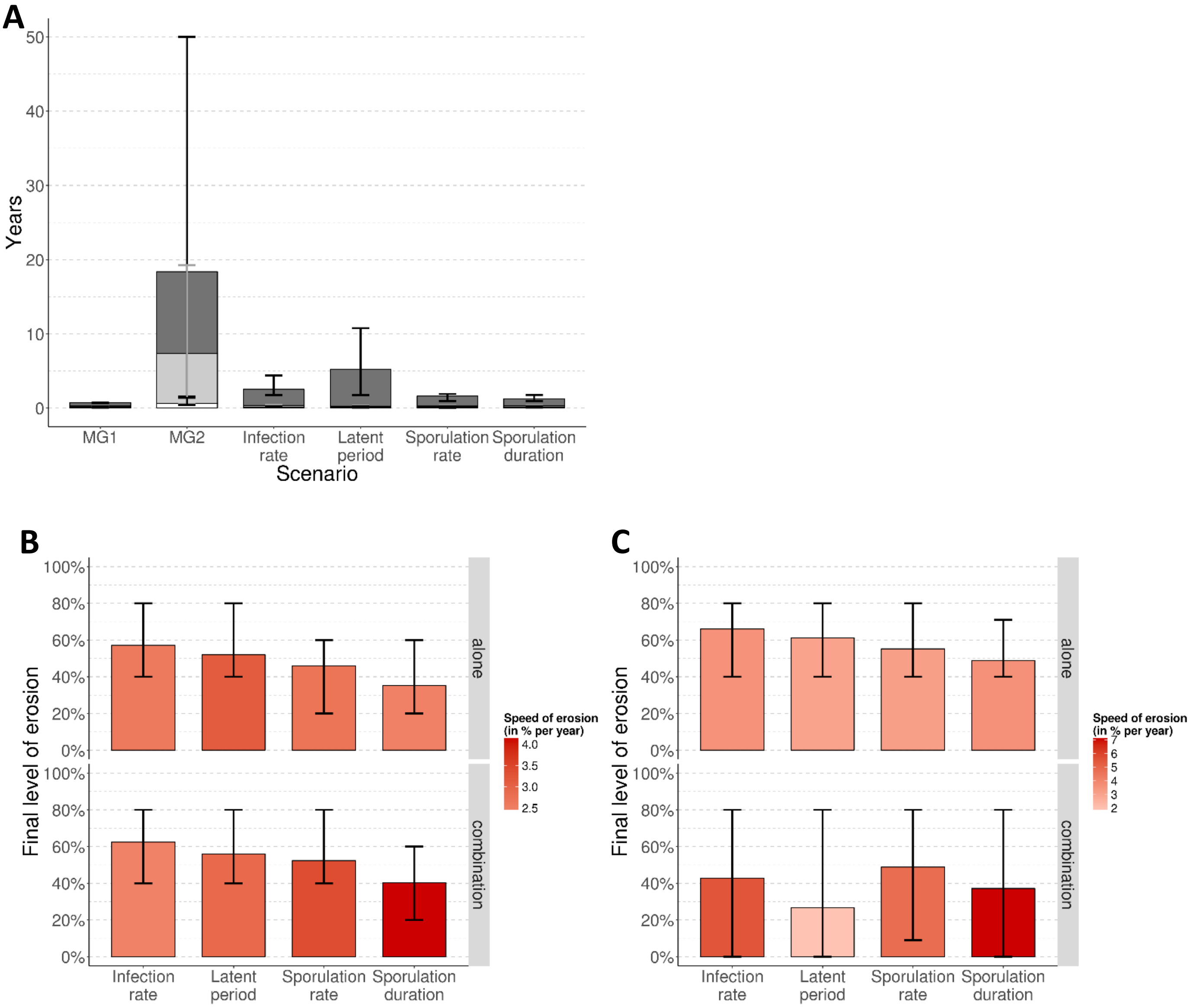


**S3 Figure. Evolutionary outcome when quantitative resistance is highly effective.** Here, the landscape is composed of a susceptible cultivar, and a resistant cultivar (cropping ratio: 80%) carrying a major gene (MG, efficiency 100%) or a quantitative resistant trait (efficiency 80% in A and B; 90% in C), alone or in combination. (A) Durability of the major gene: time to appearance of mutants (white), to first infection (light grey) or to establishment (dark grey) on resistant hosts. (B-C) Final level of erosion of quantitative resistance traits when deployed alone (top) or in combination with a major gene (bottom). The red shading indicates the average speed of erosion. Every scenario is replicated 50 times. Vertical lines show the 90% central range.

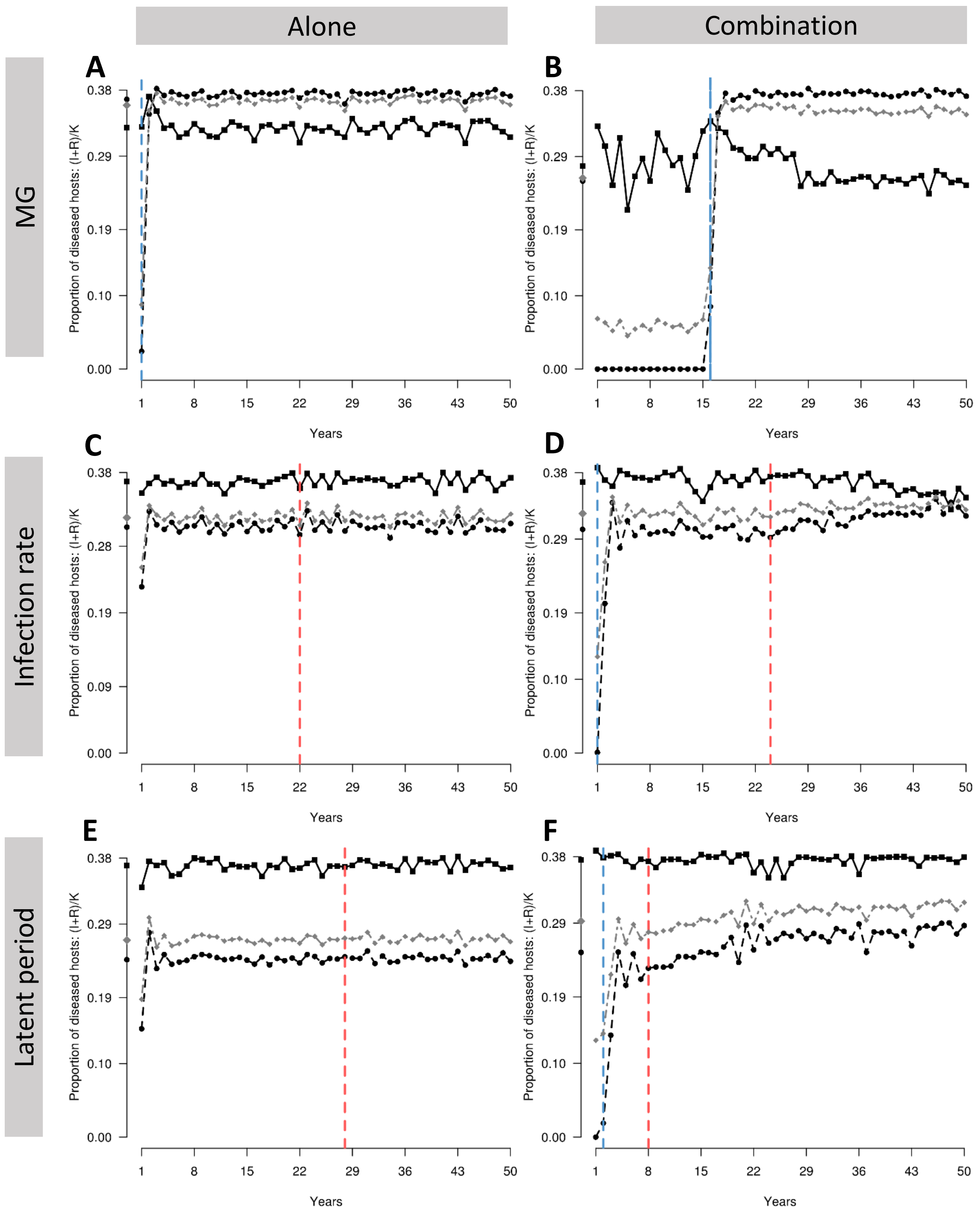


(see legend on next page)

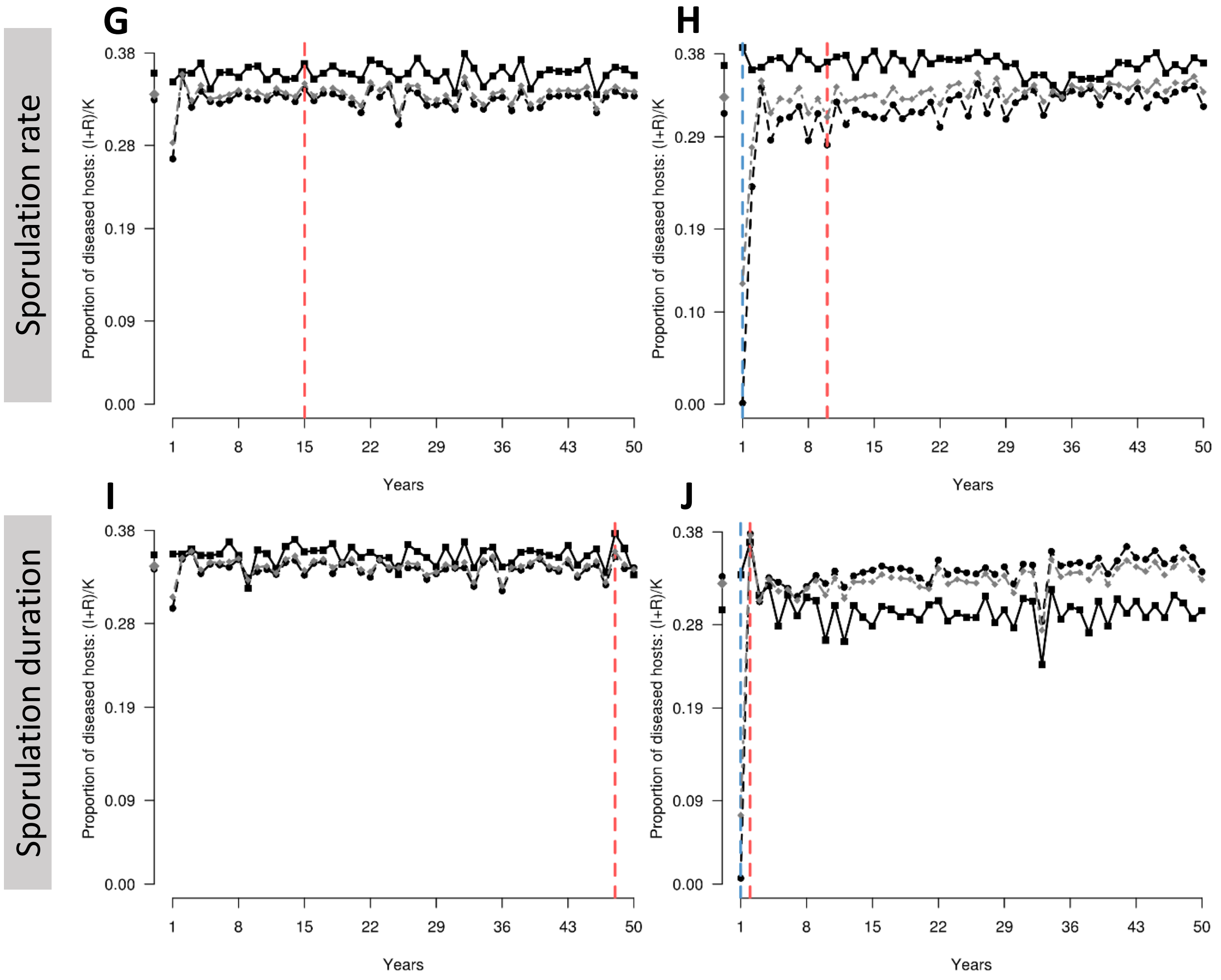


**S4 Figure. Dynamics of diseased hosts from different scenarios of resistance deployment.** Solid black curve: susceptible cultivar. Dashed black curve: resistant cultivar, carrying a major gene (A, B) or quantitative resistance against the infection rate (C, D), the latent period duration (E, F), the sporulation rate (G, H) or the sporulation duration (I, J), alone (A, C, E, G, I) or in combination with a major gene (B, D, F, H, J). Grey curve: whole landscape. The vertical blue lines mark the times to major gene breakdown; the vertical red lines mark the beginning of quantitative resistance erosion. The average AUDPC in a 100%-susceptible landscape is 0.38.

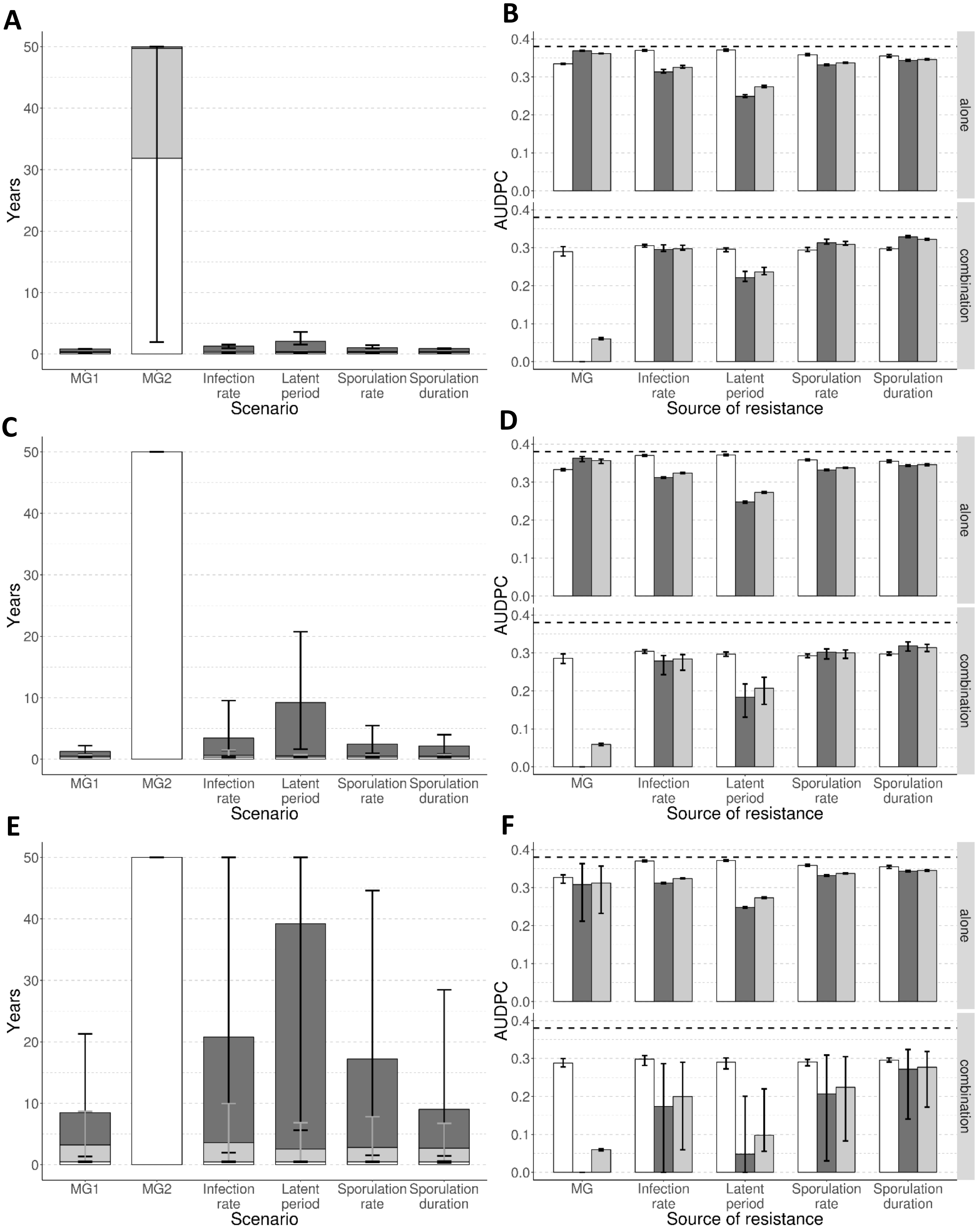


**S5 Figure. Evolutionary and epidemiological outputs when the mutation probabilities are smaller.** Here, the landscape is composed of a susceptible cultivar, and a resistant cultivar (cropping ratio: 80%) carrying a major gene (MG, efficiency 100%) or a quantitative resistant trait (efficiency 50%), alone or in combination. (A, C, E) Durability of the major gene: time to appearance of mutants (white), to first infection (light grey) or to establishment (dark grey) on resistant hosts. (B, D, F) Bars indicate the average area under disease progress curve (AUDPC) of the susceptible (white) and the resistant (dark grey) cultivars, as well as the whole landscape (light grey). The horizontal dashed line represents the average AUDPC in a fully susceptible landscape. Every scenario is replicated 50 times. Vertical lines show the 90% central range. Mutation probabilities have been set to $\tau_g=\tau_w=10^{-5}$ (A-B); $10^{-6}$ (C-D); $10^{-7}$ (E-F).

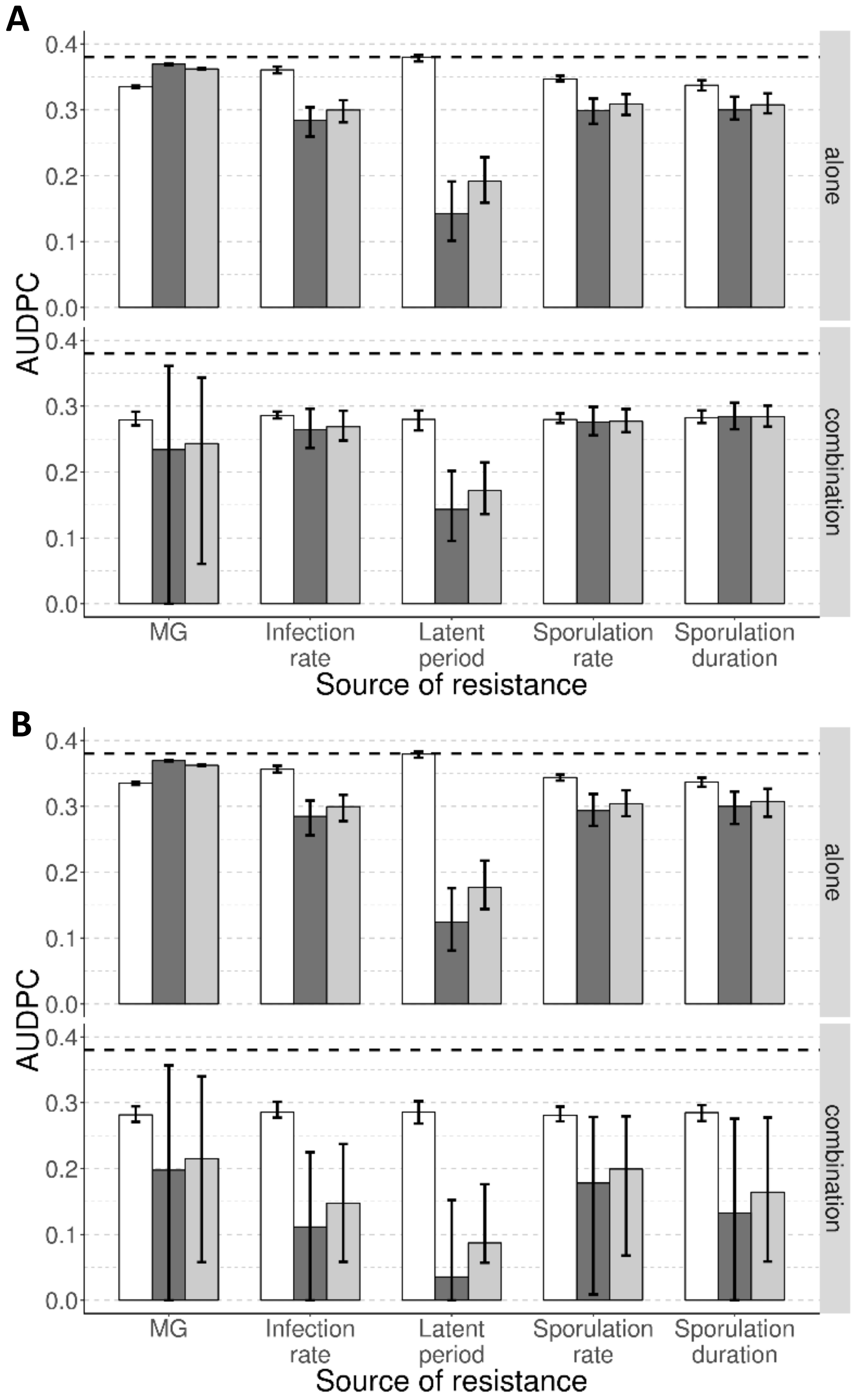


**S6 Figure. Epidemiological outcome when quantitative resistance is highly effective.** Here, the landscape is composed of a susceptible cultivar, and a resistant cultivar (cropping ratio: 80%) carrying a major gene (MG, efficiency 100%) or a quantitative resistant trait (efficiency 80% in A; 90% in B), alone (top) or in combination (bottom). Bars indicate the average area under disease progress curve (AUDPC) of the susceptible (white) and the resistant (dark grey) cultivars, as well as the whole landscape (light grey). The horizontal dashed line represents the average AUDPC in a fully susceptible landscape. Every scenario is replicated 50 times. Vertical lines show the 90% central range.

### S1 Text. Model parameterisation to rust diseases

This section details the parameterisation of the model to approximate biotrophic foliar fungal diseases as typified by rusts of wheat, caused by fungi of the genus *Puccinia* (without targeting any specific rust species). Cited references are listed in the main text.

#### Pathogen aggressiveness components

**Duration of latent and sporulation periods**. We found 36 estimates from 29 studies of different rust pathogens which included information on the duration of the latent period; similarly, there were 18 estimates from 11 studies with data on the sporulation period. All of these estimates were derived from fully susceptible hosts grown in controlled conditions (S1 Table). Standardised conditions in growth chambers were preferred over field data to facilitate comparison between studies. When available, we selected estimates in which latent periods were measured as the time from inoculation until 50% of the total number of lesions began sporulating (the most robust measure of the latent period [75-77]), otherwise the time until appearance of the first sporulating lesion was used. A Gamma distribution was fit to the data using a maximum likelihood approach (S7 Figure). From this analysis, the parameters associated with latent and sporulation periods were estimated as: $\gamma_{min}$=10; $\gamma_{var}$=9; and $\Upsilon_{max}$=24; $\Upsilon_{var}$=105.

**Infection rate**. In an experiment on the dispersal of wheat brown rust, the probability that a propagule deposited on a leaf triggers an infection was estimated to lie between 0 and 0.4, depending on the propensity of the leaf to be infected [78]. In the present study, given that the propensity of a leaf to be infected is explicitly modelled by a sigmoid contamination function, the maximal expected infection rate was set at $e_{max}$=0.4.

**Sporulation rate**. The real number of spores produced daily by a lesion and effectively dispersed to a leaf where they may trigger an infection, often summarised by 'effective sporulation rate' and denoted as $r_{max}$ here, is extremely complicated to estimate. Nevertheless, we found one study [79] where a field experiment on the spread of *P. striiformis* enabled estimation of $e_{max} \times r_{max}$ at 5 infections.day$^{-1}$. This would suggest that $r_{max}$ should be 12.5 spores.day$^{-1}$. However, in our application case, this would represent an extremely aggressive pathogen. As an indication, the basic reproductive number for such a pathogen (i.e. the theoretical number of secondary infections from a single infectious host [80]) would be $R_{0\,max} = e_{max} \times \Upsilon_{max} \times r_{max} = 120$. Recently, the $R_0$ of *P. striiformis* in large fields has been estimated to be of the order of 30 [81]. Thus, in order to simulate a reasonably aggressive pathogen, and since very few data were available to quantify $r_{max}$ compared to other parameters, $r_{max}$ has been adjusted to be 3.125 spores.day$^{-1}$, thus $R_{0\,max} = 30$.

**S1 Table. Available data on the duration of latent and sporulation periods for rust diseases caused by fungi of the genus *Puccinia*.** In all these studies, both latent and sporulation periods were determined on fully susceptible hosts under controlled conditions. Different methods to estimate duration of the latent period were used by various authors: the time from inoculation until the appearance of the first sporulating lesion (red cells), or until 50% of the lesions began sporulating (green cells). When the method is unspecified, the background of the cell is left blank.

| Reference | *Organism* | Disease | Latent period (days) | | | Sporulation period (days) | | |
|---|---|---|---|---|---|---|---|---|
| | | | min | max | mean | min | max | mean |
| [82] | *P. coronata* | Oat crown rust | 6.00 | 10.00 | 8.00 | | | |
| [83] | *P. coronata* | Oat crown rust | 6.00 | 10.00 | 8.00 | 11.0 | 16.0 | 13.5 |
| [83] | *P. graminis* | Oat stem rust | 6.00 | 10.00 | 8.00 | 17.0 | 24.0 | 20.5 |
| [84] | *P. graminis* | Wheat stem rust | 12.25 | 7.25 | 9.75 | | | |
| [85] | *P. graminis* | Wheat stem rust | 8.00 | 14.00 | 11.00 | | | |
| [83] | *P. graminis* | Wheat stem rust | 6.00 | 10.00 | 8.00 | 17.0 | 24.0 | 20.5 |
| [86] | *P. hordei* | Barley leaf rust | | | 5.00 | | | |
| [83] | *P. hordei* | Barley leaf rust | 6.00 | 10.00 | 8.00 | 17.0 | 24.0 | 20.5 |
| [87] | *P. hordei* | Barley leaf rust | 5.30 | 13.50 | 9.40 | 14.1 | 29.5 | 21.8 |
| [83] | *P. recondita* | Triticale brown rust | 6.00 | 10.00 | 8.00 | 11.0 | 16.0 | 13.5 |
| [88] | *P. recondita* | Wheat brown rust | 6.00 | 10.00 | 8.00 | 52.0 | 64.0 | 58.0 |
| [89] | *P. recondita* | Wheat brown rust | | | 8.00 | | | 21.0 |
| [83] | *P. recondita* | Wheat brown rust | 6.00 | 10.00 | 8.00 | 17.0 | 24.0 | 20.5 |
| [90] | *P. recondita* | Wheat brown rust | | | 8.00 | 20.0 | 27.0 | 23.5 |
| [91] | *P. recondita* | Wheat brown rust | 8.30 | 10.10 | 9.20 | 17.8 | 33.9 | 25.9 |
| [92] | *P. striiformis* | Barley stripe rust | 12.00 | 14.00 | 13.00 | | | |
| [93] | *P. striiformis* | Barley stripe rust | 9.00 | 15.70 | 12.35 | | | |
| [94] | *P. striiformis* | Wheat stripe rust | 12.20 | 13.95 | 13.08 | | | |
| [95] | *P. striiformis* | Wheat stripe rust | 12.60 | 13.10 | 12.85 | | | |
| [96] | *P. striiformis* | Wheat stripe rust | 12.03 | 14.27 | 13.15 | | | |
| [97] | *P. striiformis* | Wheat stripe rust | 13.00 | 15.10 | 14.05 | | | |
| [98] | *P. striiformis* | Wheat stripe rust | 10.00 | 24.00 | 17.00 | 9.0 | 13.0 | 11.0 |
| [99] | *P. striiformis* | Wheat stripe rust | 6.10 | 13.00 | 9.55 | | | |
| [100] | *P. striiformis* | Wheat stripe rust | 11.40 | 15.10 | 13.25 | | | |
| [101] | *P. striiformis* | Wheat stripe rust | | | 11.20 | | | |
| [102] | *P. striiformis* | Wheat stripe rust | 12.40 | 19.20 | 15.80 | | | |
| [89] | *P. striiformis* | Wheat stripe rust | | | 10.00 | | >29 | 29.0 |
| [83] | *P. striiformis* | Wheat stripe rust | 6.00 | 10.00 | 8.00 | 17.0 | 24.0 | 20.5 |
| [79] | *P. striiformis* | Wheat stripe rust | | | 17.00 | | | 14.0 |
| [103] | *P. striiformis* | Wheat stripe rust | 14.80 | 19.20 | 17.00 | | | |
| [104] | *P. triticina* | Wheat leaf rust | 8.20 | 9.30 | 8.75 | | | |
| [105] | *P. triticina* | Wheat leaf rust | 7.00 | 10.00 | 8.50 | | | |
| [106] | *P. triticina* | Wheat leaf rust | | | 7.75 | | | |
| [107] | *P. triticina* | Wheat leaf rust | 6.00 | 8.00 | 7.00 | 20.0 | 30.0 | 25.0 |
| [108] | *P. triticina* | Wheat leaf rust | | | 9.00 | | | 30.0 |

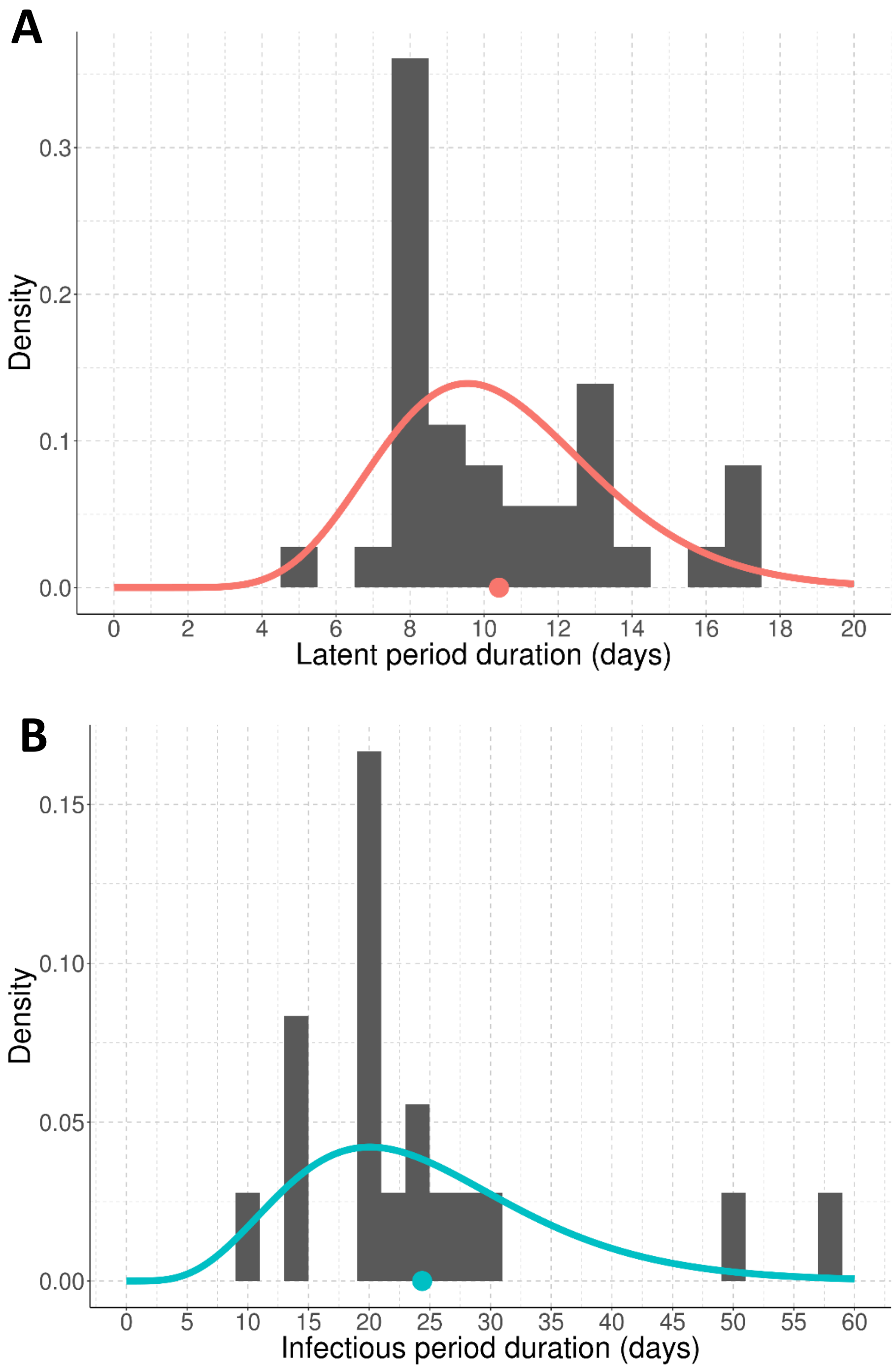


**S7 Figure. Distribution of the latent period duration (A) and length of the sporulation period (B) of rust diseases caused by fungi of the genus *Puccinia.*** Raw data were obtained from studies estimating these parameters on fully susceptible cereal cultivars (see S1 Table). Red and blue curves are Gamma distributions estimated from the data through maximum of likelihood (function *fitdistr* of the R package MASS, v7.3-47); dots indicate the mean latent period (red, 10.41 days) and sporulation period (blue, 24.37 days). The variance is 8.80 days for the latent period and 105.12 days for the sporulation period.

#### Pathogen dispersal

Based on the results of previous studies designed to estimate the dispersal kernels of *P. lagenophorae* [109] and *P. striiformis* [81, 110], the power-law function has a good ability to predict the dispersal of rust spores, assuming it is isotropic (i.e. uniform in all directions). We consequently use this function in our model:

$$g(\|z' - z\|) = \frac{(b-2)(b-1)}{2.\pi.a^2}.\left(1 + \frac{\|z'-z\|}{a}\right)^{-b} \quad (23)$$

where a > 0 is a scale parameter and b > 2 determines the weight of the dispersal tail. The expected dispersal distance is given by: $\mu_{exp} = \frac{2a}{(b-3)}$.

However, the estimation of parameters a and b is not straightforward. Indeed, many studies dealing with the dispersal of rust spores did not use a power-law function [78, 111], or they did but used a unidimensional or modified power-law functions [81, 109, 110, 112, 113], which makes calibration of our own function difficult. We therefore focused on the results obtained in four studies [78, 109, 110, 113] which used exponential functions as dispersal kernels. In spite of a lower goodness of fit of the exponential function to model spore dispersal [81, 110, 113], estimation of a mean dispersal distance from this function is easier and enables direct comparison between studies. The overall mean dispersal distance in these four studies is 19.46 m (S2 Table), thus we calibrated a and b of our power-law function in such a way that $\mu_{exp}$=20 m, which represents 1% of landscape length. Among the possible solutions we arbitrarily chose a=40 and b=7 (S8 Figure).

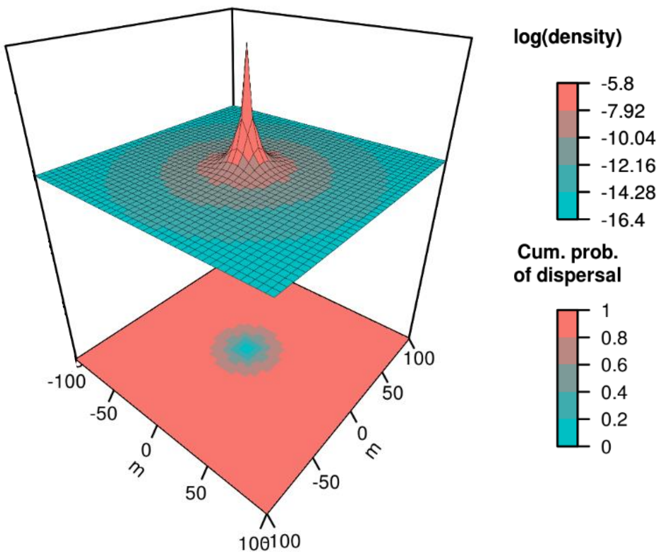


**S8 Figure. Two-dimensional representation of the power-law dispersal kernel used in this study ($\mu_{exp}$=20 m; a=40; b=7.).** The top panel indicates the logarithm of the probability to disperse from the origin to any point of the landscape, and the bottom panel indicates the cumulative probability to disperse over a given distance.

**S2 Table. Mean dispersal distance of rust spores computed from available results obtained in studies which used exponential functions.** Let the dispersal function be written as $f(x) = b.e^{-\frac{x}{a}}$ with x the distance of dispersal. Then the mean dispersal distance is simply $\mu_0 = 2.a$.

| Reference | Pathogen | Mean dispersal distance (m) |
|---|---|---|
| [113] | *P. graminis* | 36.50 |
| | *P. graminis* | 8.51 |
| | *P. graminis* | 94.76 |
| | *P. polysora* | 18.18 |
| | *P. polysora* | 11.11 |
| | *P. sorghi* | 2.22 |
| [109] | *P. lagenophorae* | 0.40 |
| [110] | *P. striiformis* | 18.87 |
| | *P. striiformis* | 1.99 |
| | *P. striiformis* | 13.42 |
| | *P. striiformis* | 8.16 |
| | *P. striiformis* | 3.04 |
| | *P. striiformis* | 1.91 |
| | *P. striiformis* | 1.98 |
| | *P. striiformis* | 0.66 |
| [78] | *P. recondita* | 38.60 |
| | *P. recondita* | 34.00 |
| | *P. recondita* | 27.80 |
| | *P. recondita* | 38.00 |
| | *P. recondita* | 29.00 |
| | **Overall mean** | **19.46 m** |

The parameters of the sigmoid curve for contamination of hosts by propagules (κ=5.33, σ=3) have been parameterised as in previous works [40, 42, 55], such as π(0)=0, π(1)=1, and the inflexion point is located at $x_0 = ((\sigma - 1)/\kappa\sigma)^{1/\sigma} \approx 0.5$. Thus, the contamination of a healthy host is easier when the proportion of healthy hosts is higher than 50%, and harder otherwise (S9 Figure).

#### Initial conditions and seasonality

The host dynamics (host growth, $\delta_v$ = 0.1 day$^{-1}$; plantation density, $C_v^0$ = 0.1 m$^{-2}$; maximal density, $C_v^{max}$ = 2 m$^{-2}$) was parameterised as in previous works [45], in order to obtain reasonable host dynamics in the absence of disease ($GLA_{TOT}$=1.48 day$^{-1}$.m$^{-2}$, noting that host dynamics is deterministic, S10 Figure).

The probability of infection at the beginning of the simulation (ϕ=5.10$^{-4}$) and the off-season survival probability (λ=10$^{-4}$) have been set such that epidemics in a fully susceptible landscape are regular (average of 10 simulations: $GLA_{TOT}$=0.48 day$^{-1}$.m$^{-2}$ [standard deviation: 0.005], $AUDPC_{TOT}$=0.38 [0.001], see the output of one simulation in S11 Figure).

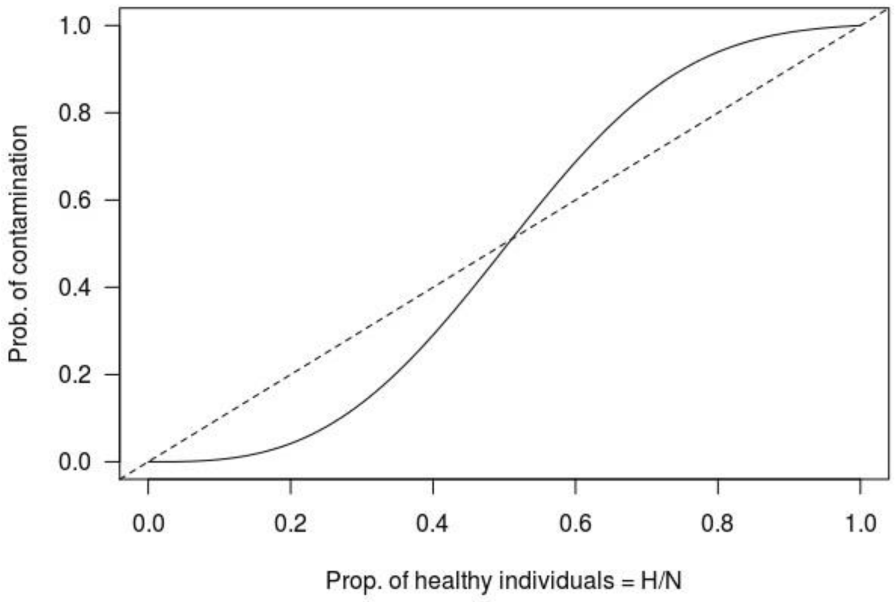


**S9 Figure. Sigmoid contamination function.** Probability for a healthy host to be contaminated following propagule arrival. The equation is $\pi(x) = \frac{1-e^{-\kappa.x^{\sigma}}}{1-e^{-\kappa}}$ with κ=5.33 and σ=3.

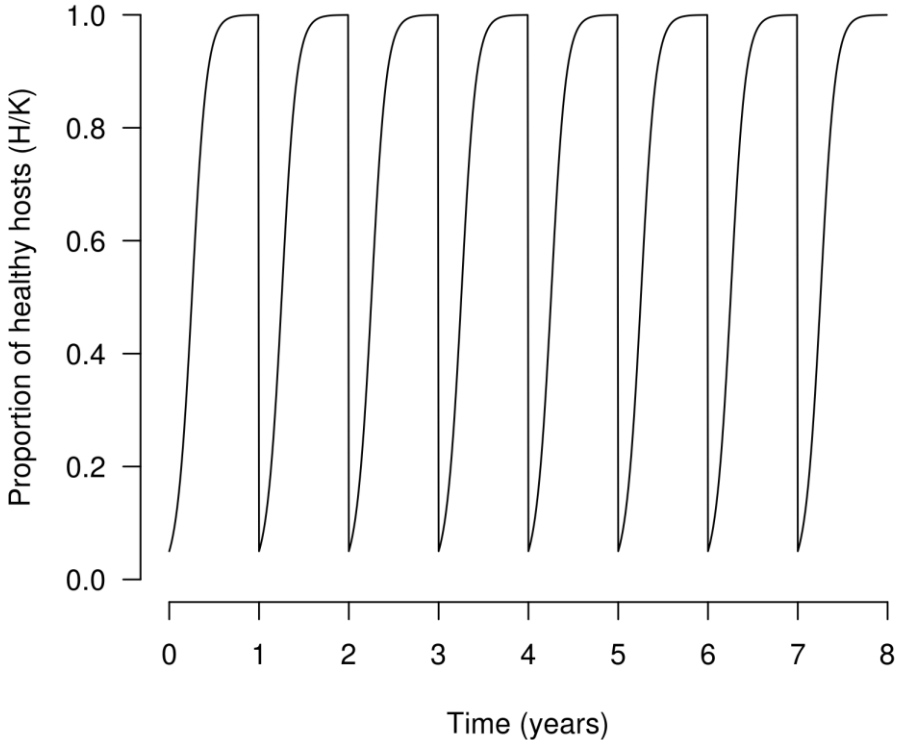


**S10 Figure. Deterministic dynamics of healthy hosts in the absence of disease.**
$GLA_{TOT}$=1.48 $day^{-1}.m^{-2}$.

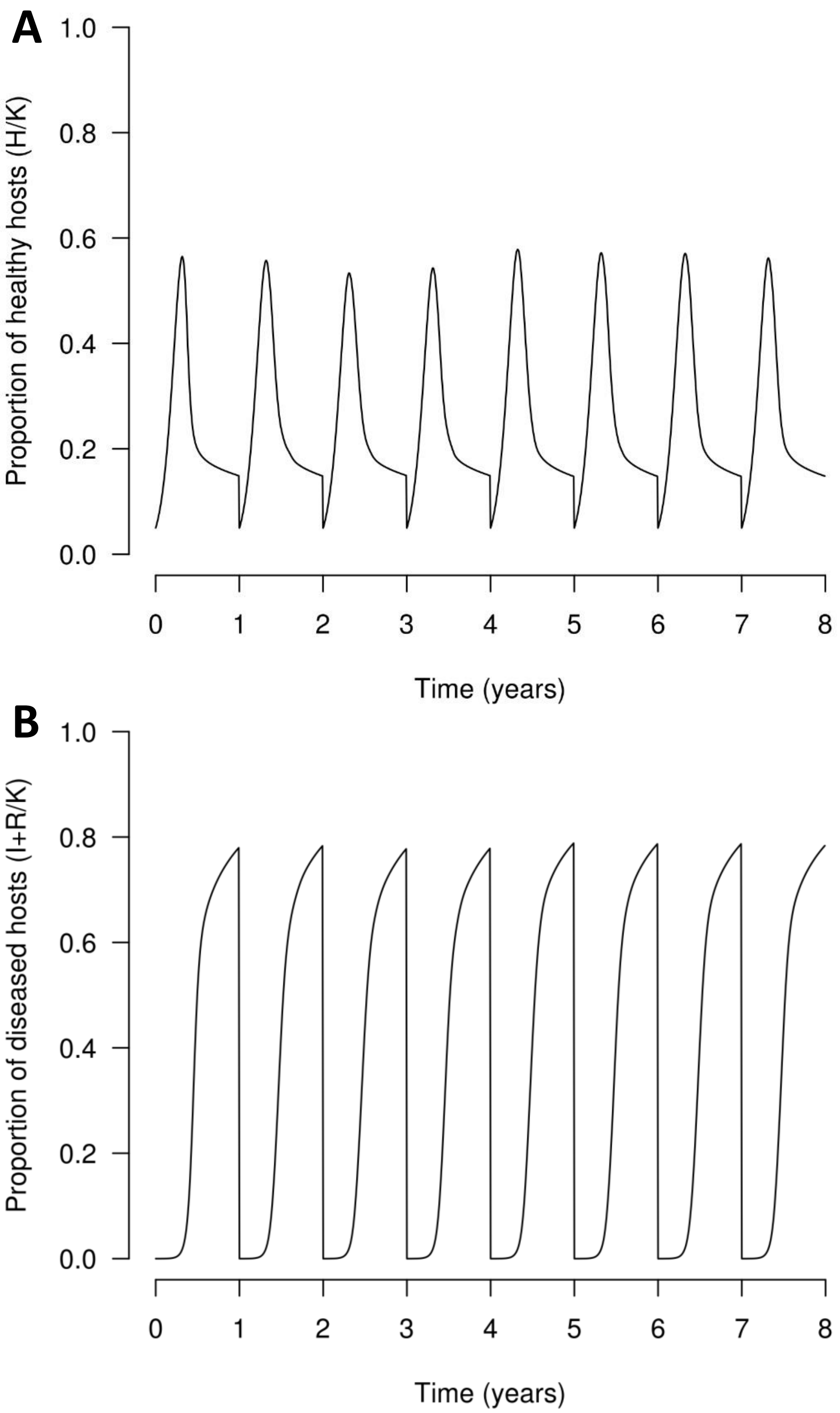


**S11 Figure. Dynamics of healthy** (A, $GLA_{TOT}$=0.49 day$^{-1}$.m$^{-2}$) **and diseased** (B, $AUDPC_{TOT}$=0.38) **hosts in a fully susceptible landscape in a simulated example.**

#### Evolutionary parameters

Very few empirical data are available from rust pathogens to help parameterise the model with respect to mutation probabilities, cost of infectivity, cost of aggressiveness and number of mutations required to completely erode a trait for quantitative resistance. It is important to remember that the mutation probability in the model refers to the probability that a spore has a different phenotype from its parental lesion. For instance, $\tau_1$ gives the probability for a spore to have an infective phenotype on a resistant cultivar carrying major gene 1, given that this spore was produced by a lesion triggered by a non-infective pathogen. This probability depends on the number of mutations per generation per base pair (i.e. the classic 'genetic mutation rate' of empirical studies), the number and nature of the specific genetic mutations required to overcome major gene 1, and the potential dependency between these mutations. In addition to the lack of empirical data, all these parameters are likely to vary depending on the pathosystem as well as the resistance sources.

Consequently, this work focuses on a simple theoretical case, where evolutionary parameters have been arbitrarily fixed. Since one of the objectives of this application case is to identify promising combinations of qualitative and quantitative resistance, and based on preliminary simulations, the mutation probabilities were set at $\tau_g=\tau_w=10^{-4}$ in such way that a cultivar carrying a single major gene would be overcome in less than one year. Using these values, pathogen evolution is quick enough to potentially adapt to the resistance genes, but slow enough to allow comparison of the effectiveness of different resistance combinations with regard to controlling disease. The resistance efficiency is set at $\rho_g=1$ for qualitative resistance (i.e. a major resistance gene confers complete immunity against non-adapted pathogens), and $\rho_w=0.5$ for quantitative resistance traits (i.e. infection rate, sporulation rate, duration of the sporulation period, or number of epidemic cycles in a cropping season of a non-adapted pathogen on resistant hosts is reduced by 50%). The cost of infectivity and the cost of aggressiveness were set at $\theta_g=\theta_w=0.5$, the trade-off strength at $\beta_w=1$ (linear trade-off), and the number of pathotypes with regard to the aggressiveness components at $Q_w=6$ (i.e. every mutation step improves or degrades an aggressiveness component by 20%). This last value is a compromise between the high number of steps required to simulate a gradual pathogen evolution, and having a smaller number of different pathotypes to limit the computational time required to perform the simulations.

This parameterisation gives the following infectivity and aggressiveness matrices:

| **Infectivity matrix for major gene g** INF$^g$ | | **Host genotype v**<br>Susceptible (SC) $mg_g(v)=0$ | **Host genotype v**<br>Resistant (RC) $mg_g(v)=1$ |
|---|---|---|---|
| **Pathogen genotype p** | Non-infective $ig_g(p)=0$ | 1 | 0 |
| **Pathogen genotype p** | Infective $ig_g(p)=1$ | 0.5 | 1 |

| **Aggressiveness matrix for component w** AGG$^w$ | | **Host genotype v**<br>Susceptible (SC) $qr_w(v)=0$ | **Host genotype v**<br>Resistant (RC) $qr_w(v)=1$ |
|---|---|---|---|
| **Pathogen genotype p** | Non-aggressive $ag_w(p)=1$ | 1 | 0.5 |
| **Pathogen genotype p** | $ag_w(p)=2$ | 0.9 | 0.6 |
| **Pathogen genotype p** | $ag_w(p)=3$ | 0.8 | 0.7 |
| **Pathogen genotype p** | $ag_w(p)=4$ | 0.7 | 0.8 |
| **Pathogen genotype p** | $ag_w(p)=5$ | 0.6 | 0.9 |
| **Pathogen genotype p** | Fully aggressive $ag_w(p)=6$ | 0.5 | 1 |

Grey lines indicate pathogen population at the beginning of the simulations.

The parameterisation of the mutation probabilities ($\tau_g=10^{-4}$ and $\tau_w=10^{-4}$) and the number of steps to completely erode a quantitative resistance (given by $Q_w$) give the following mutation matrix for infectivity gene g:

| | $ig_g(p)=0$ | $ig_g(p)=1$ |
|---|---|---|
| $ig_g(p)=0$ | 0.9999 | $10^{-4}$ |
| $ig_g(p)=1$ | $10^{-4}$ | 0.9999 |

and the following mutation matrix for aggressiveness component w:

| | $ag_w(p)=1$ | $ag_w(p)=2$ | $ag_w(p)=3$ | $ag_w(p)=4$ | $ag_w(p)=5$ | $ag_w(p)=6$ |
|---|---|---|---|---|---|---|
| $ag_w(p)=1$ | 0.9999 | $10^{-4}$ | 0 | 0 | 0 | 0 |
| $ag_w(p)=2$ | $5.10^{-5}$ | 0.9999 | $5.10^{-5}$ | 0 | 0 | 0 |
| $ag_w(p)=3$ | 0 | $5.10^{-5}$ | 0.9999 | $5.10^{-5}$ | 0 | 0 |
| $ag_w(p)=4$ | 0 | 0 | $5.10^{-5}$ | 0.9999 | $5.10^{-5}$ | 0 |
| $ag_w(p)=5$ | 0 | 0 | 0 | $5.10^{-5}$ | 0.9999 | $5.10^{-5}$ |
| $ag_w(p)=6$ | 0 | 0 | 0 | 0 | $10^{-4}$ | 0.9999 |

#### Discussion of model parameterisation and assumptions

##### Parameterisation to rust pathogens.

**Pathogen aggressiveness.** This model has been calibrated for rust-like pathogens using available knowledge of the epidemiology of fungi in the genus *Puccinia*. In particular, we found many data related to pathogen aggressiveness and dispersal. This allows the reliable estimates of parameters $e_{max}$, $\gamma_{min}$, $\gamma_{var}$, $\Upsilon_{max}$ and $\Upsilon_{var}$. However, as noted above, the calculation of the effective number of spores produced daily by a lesion and effectively dispersed to a leaf where they may trigger an infection ($r_{max}$ in our model) is extremely challenging. Acquisition of data to inform this parameter is crucial, since this parameter has a strong influence on epidemic spread and pathogen evolution [45]. In this study, $r_{max}$ directly affects pathogen population size, which influences both the number of new infected hosts from a single infectious source, and the probability that some of these new infections are triggered by mutant spores. For example, smaller values should lead to higher times to appearance of mutants (durability measure (d1)) than what we observed in our simulations (see white bars in Fig 5A).

**Pathogen dispersal.** The dispersal of fungal spores has been extensively studied, and we now have considerable evidence to support the use of a power-law kernel to simulate spore dispersal [81, 109, 110]. Available data indicate that the mean dispersal distance is of the order of 20 m (see S2 Table). However, sensitivity analyses of models simulating plant epidemics show that both the mean dispersal distance [40, 114] and the width of the tail of the dispersal kernel [42, 79] have a great impact on pathogen spread. In the present work, the calibration of parameters a and b gives an appropriate mean dispersal distance, but is probably less accurate with respect to simulation of long-distance dispersal events (determined by the width of the tail of the dispersal kernel). In the future, we need studies built with exactly the same equations, to better compare and calibrate parameter values of dispersal kernels. This will be particularly important in future modelling studies aiming to explore the effect of landscape composition (e.g. proportion and spatial aggregation of a resistant cultivar) on pathogen spread and evolution (see 'Future research directions' in the discussion section).

All healthy sites of a plant do not have the same propensity to be infected [78]. This is at least partly due to plant architecture, which makes healthy sites not equally accessible to spores following dispersal. Thus, once the most accessible sites are infected, the remaining healthy sites may be harder to reach and infect. Our contamination function (see S9 Figure) accounts for this process. The high flexibility of this function makes it relevant for a wide range of pathogens, including rusts. However, we did not have any experimental data to parameterise it for cereal crops. Therefore, we used the same arbitrary values (κ, σ) as in previous works [40, 55] in order to facilitate comparisons between studies. Nevertheless, since this function only mitigates the overall infection rate ($e_{max}$), it is likely that epidemic spread depends more on the value of the latter than on the parameters or shape of this sigmoid function. A sensitivity analysis of a similar simulation model showed that use of a linear contamination function, rather than a sigmoid function [41] has little effect on the level of pathogen adaptation to quantitative resistance.

**Seasonality and initial conditions.** Parameters related to host growth ($\delta_v$), host density ($C_v^{max}$), initial conditions ($C_v^0$ and ϕ), and seasonality (λ) are specific to the agricultural (e.g. choice of cultivars, planting density) and epidemiological (e.g. initial level of contamination, presence of a wild reservoir or volunteer host plants for the pathogen) contexts. In our application case, these parameters have been parameterised so as to simulate regular host dynamic and epidemics. This helps compare different resistance deployment strategies in standardised conditions, and attribute variation in model outputs to the deployment strategies rather than to the initial conditions or the epidemiological context. On the other hand, it could be interesting to vary the growth rate of resistant cultivars (or their contribution to yield) to study the impact of cost of resistance on the performance of different deployment strategies. One could also vary the off-season survival probability of infectious hosts (λ) to investigate the potential of strategies based on green bridge suppression (e.g. fungicide treatments in the end of the cropping season; removal of potential volunteer plants during the off-season).

**Pathogen evolution.** As discussed previously, parameterisation of evolutionary processes in the model from empirical data is extremely difficult. In addition to the scarcity of studies designed to estimate such parameters for rusts, these parameters may not correspond with the definition of the parameters in our model (e.g. our 'mutation probability' is different from the classic 'mutation rate'; see above). Consequently, we chose to investigate combinations of qualitative and quantitative resistances in a simple context, using the same mutation probabilities ($\tau_g$ and $\tau_w$), costs of infectivity ($\theta_g$) and aggressiveness ($\theta_w$), trade-off strengths ($\beta_w$), and numbers of steps to completely erode a quantitative resistant trait ($Q_w$) for every major resistance gene g and trait of quantitative resistance w.

In this model, the mutation probability gives the probability of appearance of mutants. Therefore it has a great effect on the time to appearance of such mutants [30, 39] or the speed of erosion of quantitative resistance [56]. However, provided that mutation probabilities are the same for all infectivity genes and aggressiveness components (i.e. $\tau_g = \tau_w = constant, \forall g, \forall w$), the specific value of these mutation probabilities should similarly affect all scenarios of our numerical experimental design. In other words, they should not change the relative durability and epidemiological efficiency of these simulated scenarios, and consequently the ranking of the tested resistance combinations. To test this hypothesis, we replicated the simulations with smaller mutation probabilities: $\tau_g = \tau_w = \{10^{-5}; 10^{-6}; 10^{-7}\}$. The results of these new simulations show that the durability of the major gene increases with decreasing mutation probabilities (S5ACE Figure), but the tested resistance combinations keep the same ranking. Similarly, the combinations of qualitative and quantitative resistances result in better epidemiological outcomes when the mutation probabilities decrease (S5BDF Figure). Nevertheless, regardless the mutation probabilities, the most promising combination is still the pyramid of two major resistance genes, followed by a major gene combined with a quantitative resistance trait against the latent period, next with a quantitative resistance trait against the infection rate and the sporulation rate, and finally with a quantitative resistance trait against the sporulation duration.

In contrast, the fitness costs associated with pathogen adaptation may differentially affect the simulated scenarios. Several studies have demonstrated that higher infectivity costs increase resistance durability [29, 30] and epidemiological efficiency [33]. In our results, the durability of the pyramid of two major genes is likely due to the cost of infectivity ($\theta_g$) endured by mutant pathogens (see main text and Fig 5A). However, smaller values would result in a reduced performance of this strategy in comparison to other resistance combinations. This may also be true for comparisons of pyramiding with other spatio-temporal deployment strategies such as mosaics [50]. Similarly, the time to establishment of mutant pathogens in a host population carrying quantitative resistance has been shown to increase with higher costs of aggressiveness [37] or stronger trade-off relationships [41, 45,

55]. Thus, the associated parameters ($\theta_w$ and $\beta_w$), as well as the number of steps to erode a quantitative resistant trait ($Q_w$) should considerably impact model outputs like final level and speed of erosion of quantitative resistance.

To conclude, the impact of all these evolutionary parameters, related to the choice of the resistance source, on the evolutionary and epidemiological outcomes of the simulated deployment strategies need to be explicitly evaluated within a dedicated numerical experimental design. This is the objective of future studies (see section 'Future research directions' in main text). Furthermore, given the variability in fitness costs and trade-off relationships associated with the evolution of pathogens, including rust fungi [65], a careful assessment of these parameters is required before the deployment of genetic resistance in the field, in order to get reliable predictions of their durability and efficiency.

**Parameterisation to necrotrophic fungal pathogens**.

Parameterisation of our model is flexible enough to encompass a wide range of pathogens. As an example, although our application case focused on biotrophic fungi, necrotrophic pathogens could be simulated, by allowing transmission from stubble after crop harvest instead of from living tissue during the cropping season. To this end, the latent period could be increased to cover the whole cropping season, and the off-season survival probability could be adjusted in such a way that the total pool of produced spores is available for the following cropping season (after which the pool of spores would no longer be available). This scenario would at least partially represent major features of interactions such as that between canola and blackleg (*Leptosphaeria maculans*) [115].

#### S2 Text. Calculation of the threshold for pathogen establishment.

In this work, the time to mutant pathogen establishment in the resistant host population is defined as the point at which extinction becomes unlikely. Assessment of this time requires the calculation of a threshold (denoted by N hereafter) for the number of infections of resistant hosts by mutant pathogens, above which mutant pathogens are unlikely to go extinct.

The spread of a mutant pathogen across years from a single infection of a resistant host requires surviving the bottleneck imposed by host harvest (denoted by event $\{Surv_I\}$), and the infection of new resistant hosts at the beginning of the next cropping season (denoted by event $\{Inf_I|Surv_I\}$). Thus the probability of effective spread from this single infection (event $\{Inf_I\}$) is:

$$P(Inf_I) = P(Surv_I) \times P(Inf_I|Surv_I) \quad (1)$$

The probability for a single infection to survive the bottleneck imposed by host harvest and the off-season is simply:

$$P(Surv_I) = \lambda \quad (2)$$

We now assume that a single infection survives the bottleneck. The probability for a single spore, produced by a mutant pathogen (completely adapted to the resistant host population) to infect a resistant host (assumed to be present in the field) is given by $e_{max}$ (neglecting the probability of reverse mutation to a non-adapted pathotype, the probability to disperse outside the field, and the effect of plant architecture on the probability of contamination). Then, the probability of extinction of this single spore is $1 - e_{max}$.

Since a single infection produces a total of $r_{max} \times \Upsilon_{max}$ spores, the probability of extinction of all these spores is: $P(Ext_{Pr}) = (1 - e_{max})^{r_{max} \times \Upsilon_{max}}$. Given the values of $e_{max}$=0.40, $r_{max}$=3.125 and $\Upsilon_{max}$=24, we have $P(Ext_{Pr}) \approx 2.3 \times 10^{-17}$. Consequently, the probability of infection of new resistant hosts by a single infection is:

$$P(Inf_I|Surv_I) = 1 - P(Ext_{Pr}) \approx 1 \quad (3)$$

The combination of equations (1), (2) and (3) gives:

$$P(Inf_I) = \lambda \quad (4)$$

Then, the probability of extinction of a single infection is $P(Ext_I) = 1 - P(Inf_I) = 1 - \lambda$, and the probability of extinction of N infections is:

$$P(Ext) = P(Ext_I)^N = (1 - \lambda)^N \quad (5)$$

S12 Figure shows the probability of extinction for different values of N, with $\lambda=10^{-4}$. Above N=50,000 infections, the probability of extinction is less than 1%. Therefore, we chose this threshold to define the time to mutant pathogen establishment.

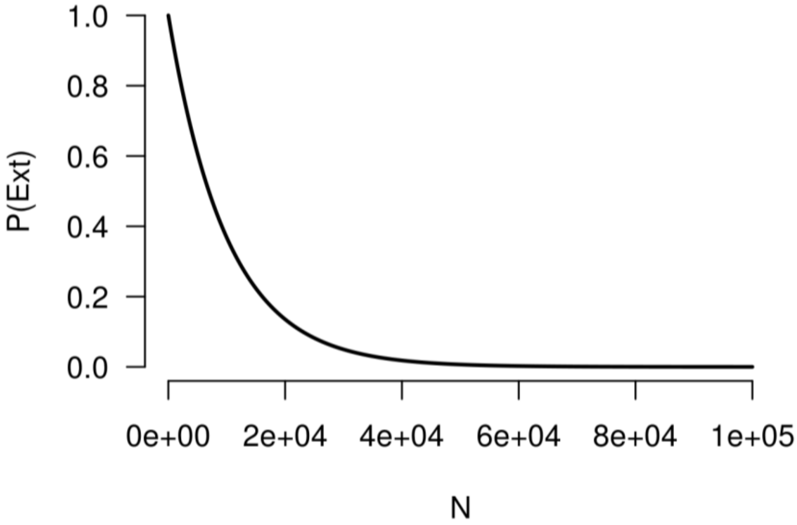


**S12 Figure. Probability of extinction of a mutant pathogen in a steady environment, depending on the number of infections.** $P(Ext) = (1 - \lambda)^N$ with $\lambda=10^{-4}$
